# High-density multi-fiber photometry for studying large-scale brain circuit dynamics

**DOI:** 10.1101/422857

**Authors:** Yaroslav Sych, Maria Chernysheva, Lazar T. Sumanovski, Fritjof Helmchen

**Affiliations:** Brain Research Institute, University of Zurich, Zurich, Switzerland; Neuroscience Center Zurich, Zurich, Switzerland

## Abstract

Animal behavior originates from neuronal activity distributed and coordinated across brain-wide networks. However, techniques to assess large-scale brain circuit dynamics in behaving animals remain limited. Here we present compact, high-density arrays of optical fibers that can be chronically implanted into the mammalian brain, enabling multi-fiber photometry as well as optogenetic perturbations across many regions. In mice engaged in a texture discrimination task we achieved simultaneous photometric calcium recordings from networks of 12 to 48 brain regions, including striatal, thalamic, hippocampal, and cortical areas. Furthermore, we optically perturbed specific subsets of regions in VGAT-ChR2 mice by using a spatial light modulator to address the respective fiber channels. Perturbation of ventral thalamic nuclei caused distributed network modulation and behavioral deficits. Finally, we demonstrate multi-fiber photometry in freely moving animals, including simultaneous recordings from two mice during social interaction. Thus, high-density multi-fiber arrays are simple, low-cost, and versatile tools that open novel ways to investigate large-scale brain dynamics during behavior.

## INTRODUCTION

Neural substrates of complex behaviors are often represented by large ensembles of neurons distributed across multiple brain areas. Whereas whole-brain imaging has become feasible at high spatiotemporal resolution in the small brains of *C. elegans*^1^, *Drosophila*^2^, and zebrafish larvae^1,3^, it remains challenging to assess distributed neural activity in the mammalian brain, especially during behavior. Available methods to address large scale dynamics are all limited in certain ways: functional magnetic resonance imaging (fMRI) infers neural activity only indirectly from blood flow changes, suffers from low spatiotemporal fidelity, and is difficult to apply to behaving animals. Functional ultrasound (fUS) imaging^4^ shows improved temporal resolution but still is primarily a measure of microvasculature dynamics. Electrophysiological approaches have traditionally been applied either to the brain surface—such as in electro-encephalogram (EEG) or electrocorticogram (ECoG) recordings—or to specific deep brain regions to measure local field potentials (LFPs) and multi- or single-unit activity. Although the latter approaches have been recently scaled up to the wider scale, for example using multisite LFP recordings^5^ or the new Neuropixels silicon probes^6^, they still face the difficulty to dissect the contribution of distinct cell types, e.g., neurons giving rise to specific long-range projections. Optical imaging approaches, combined with modern genetic tools^7^ can provide such cell-type specificity and thus nicely complement electrophysiological methods. In recent years, imaging techniques have been extended to the larger scale, too. For example, several two-photon mesoscopes have been developed, enabling cellular resolution imaging from multiple regions within millimeter-sized field of views^8–11^. However, these systems require sophisticated technology and are expensive. Wide-field camera imaging using one-photon excitation of calcium or voltage indicators in transgenic mice^12–16^ offers a simple and lower-cost alternative but sacrifices cellular resolution. Both approaches so far have been mainly applied to the brain surface. Subcortical imaging can be achieved by inserting gradient-index lenses^17^ but typically only individual deep brain regions have been targeted. It is therefore highly desirable to expand large-scale optical recordings to the broader subcortical realm where many brain nuclei exert essential functions.

A simple method to optically record from subcortical regions is fiber photometry, which collects bulk fluorescence signals excited at the tip of an optical fiber^18–20^. Although lacking cellular resolution, fiber photometry has gained considerable attention due to its technical simplicity and its versatility. Fibers can be chronically implanted deep in the brain. Furthermore, a particular strength lies in the combination with cell-type or pathway-specific expression of genetically encoded calcium or voltage indicators^20–23^. In addition, optical fibers are commonly used for optogenetic perturbations^24,25^. Fiber photometry is potentially scalable to allow mesoscopic measurements and manipulations across many brain areas. Two previous studies demonstrated fiber photometry from multiple regions in the mouse brain (with up to 7 probes across both hemispheres)^26,27^. However, because single-fiber implants with relatively bulky standard ferrules were used, the minimal distance between two neighboring fiber implants was 1.25 mm, fundamentally restricting channel density and the flexibility of implant configurations. Clearly, a higher density of channels is desirable in order to more comprehensively explore large-scale functional circuits that often contain parallel pathways^28^ and feed-forward/feed-back loops. Eventually, the goal is to obtain multi-site optical recordings with a broad coverage across the mesoscale network, containing on the order of several hundred brain regions^29^.

Here we present a major step in this direction by introducing novel high-density multi-fiber arrays that enable multi-fiber photometry from several tens of brain regions simultaneously. We built single arrays with up to 24 fiber channels and demonstrate that multiple arrays can be chronically implanted in a mouse brain in parallel and flexibly arranged to target a specific set of regions of interest. We demonstrate simultaneous measurement of calcium dynamics from up to 48 brain regions. Furthermore, we integrated a spatial light modulator in our setup to target specific regions for optogenetic circuit manipulations. We provide proof-of-concept for the combination of multi-fiber photometry with simultaneous multi-regional network perturbations. The multi-fiber arrays are novel tools that complement the existing set of large-scale recording techniques and provide new opportunities to study the functional organization and behavior-related dynamics of mesoscale circuits in the mammalian brain.

## RESULTS

### Optical setup and high-density multi-fiber arrays

We built an optical setup for fluorescence recordings through multiple optical fibers in parallel (**Fig. 1a**; Online Methods). Fibers were flexibly held together in one or several custom bundles (length 2 - 4 m) and were fixed at the back and front ends using modified fiber array connectors (US Conec; NC; Online Methods). These commercially available fiber ferrules contain guiding grooves for precise alignment of 12, 24, or 48 fibers (**Fig. 1b**; 250-µm fiber spacing). To excite fluorescence, we coupled laser light into the fiber array at the back end, guided it through the fiber bundle, and connected the front end to a front piece that was chronically implanted into the brain. In most cases, we built this front piece using a 12-fiber connector (**Fig. 1c**). In some experiments we connected separate branches of the fiber bundle to multiple front pieces, including a 24-fiber front piece (see below). We used optical fibers with 100-µm core diameter and solarization-resistant polyimide coating, which exhibits low background fluorescence and elicits minimal immune response of brain tissue (**Supplementary Figure 1**). The distal fiber ends (facing the brain) were cut flat. Their lengths can be adjusted flexibly depending on the locations of the targeted brain regions and the implantation geometry (**Fig. 1c**; Online Methods). All fibers were epoxy-fastened inside the ferrule and the proximal fiber ends were polished to achieve good optical coupling between the front piece and the matching fiber bundle connector (Online Methods). Guiding metal alloy pins and a custom housing with magnetic pins ensured precise alignment of fibers in the connector and the front piece as well as easy attachment and detachment. The dimensions of a front piece are 7-mm length, 2.45-mm width, and 8.05-mm height at a weight of 0.32 g. Fluorescence light excited at the tip of the implanted fibers was collected back through the same fiber bundles and fiber end faces were imaged onto a camera (**Fig. 1a,b**).

**Figure 1.**
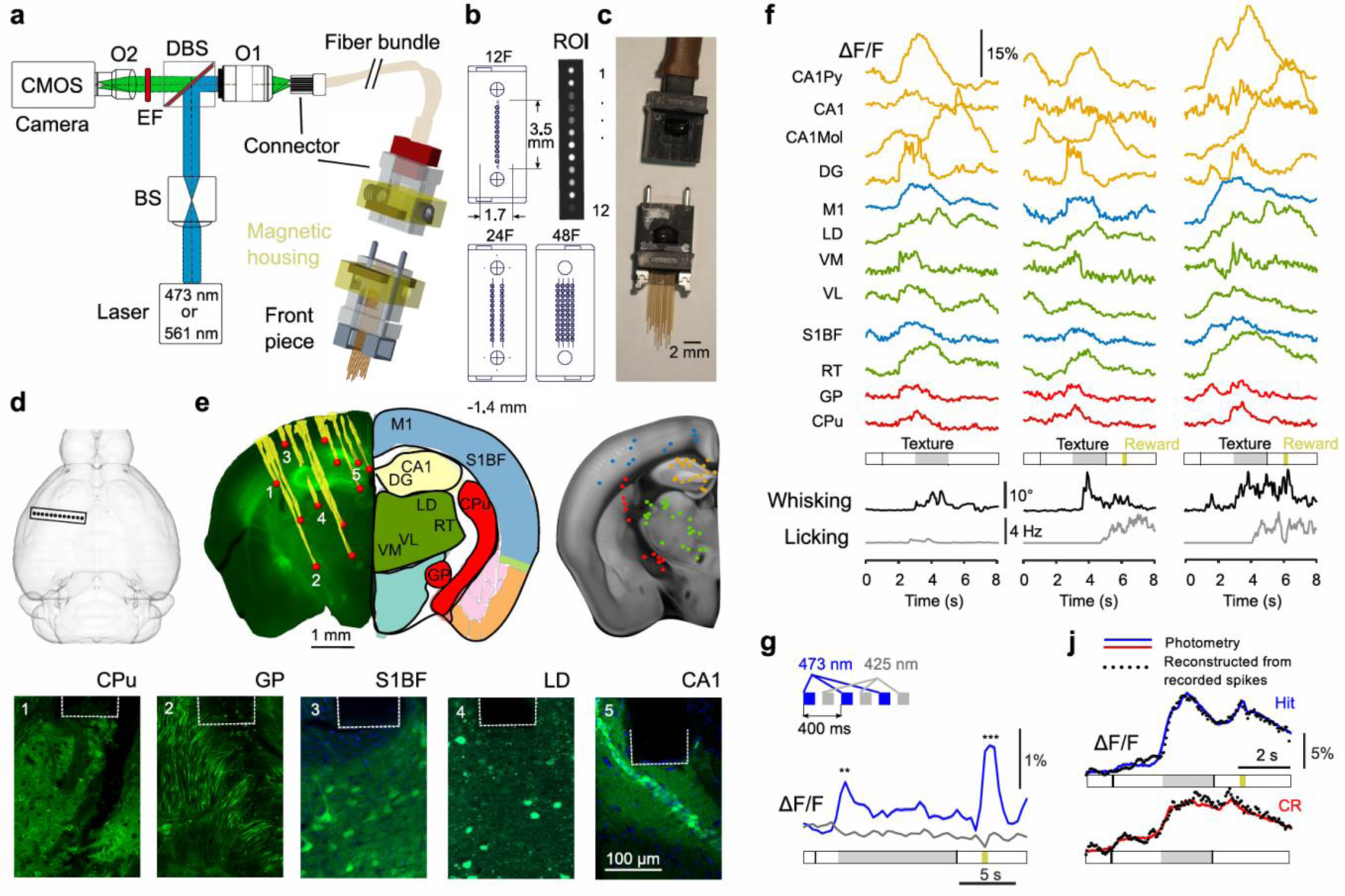
Optical setup for multi-region photometry using high-density multi-fiber arrays. (**a**) Schematic setup. The laser beam is shaped with a cylindrical lens (BS, beam shaper) to achieve a light-sheet illumination profile at the focal plane of objective 1 (O1) for coupling into the fiber bundle, which connects to the front piece. A dichroic beamsplitter (DBS) separates the fluorescence emission light from the illumination path and emission filters (EF) select the optimal wavelength range for the respective calcium indicator. Objective 2 (O2) creates an image of the fiber array on the CMOS camera sensor. We used 473-nm light to excite GCaMP6m and 561-nm light to excite R-CaMP1.07. (**b**) Dimensions of fiber ferrules holding 12, 24, and 48 fibers, respectively. Upper right inset shows an example camera image of a 12-fiber array end face with excited GCaMP6m fluorescence. (**c**) Photograph of a 12-fiber array front piece and its fiber bundle connector (without magnetic housing). (**d**) Schematic top view of a mouse brain surface and an implanted fiber array. (**e**) Histology aligned to the Allen Brain Atlas^32^. Fiber tracks were reconstructed (yellow traces) and fiber tip positions (red spheres) mapped to the corresponding brain regions. Right inset shows fiber tip positions pooled from 6 mice and colored according to the targeted brain regions (blue – cortex, red – basal ganglia, green – thalamus, yellow – hippocampus). Bottom row shows example histological confocal images displaying GCaMP6m-expressing cells below individual fiber tips. (**f**) Example calcium traces (ΔF/F) from 3 individual trials, recorded simultaneously across 12 brain regions (color code as in panel e). Whisking and licking behaviors are shown at the bottom. (**g**) Fluorescence traces (averaged over 154 trials) for two-color illumination alternating at 473 nm (blue, calcium dependent) and 425 nm (grey, calcium independent). Mean fluorescence excited at 473 nm for texture presentation onset and the lick/reward periods was significantly higher as compared to 425 nm (** p < 0.01, *** p < 0.001; one-way ANOVA). (**j**) Average photometric signals in cortical region M1 for Hit (blue: n = 112) and correct rejection CR (red: n = 120) trials from opto-tetrode control experiments. A weighted sum of single-unit spike rates measured from neurons in the vicinity of the fiber tip could well reconstruct the observed calcium signals.

We first implanted several mice (n = 6) with 12-fiber arrays targeting 12 brain regions distributed across basal ganglia, thalamus, hippocampus, and neocortex (**Fig. 1d**). All regions were previously injected with the virus expressing calcium indicator GCaMP6m^30^ (Online Methods). At the end of the experiments, we verified the 3D positions of the fiber tips from coronal brain sections, in which we could discern the fiber tracks and the labeled neuronal populations at the fiber tip locations (**Fig. 1e**; Online Methods). By exciting GCaMP6m fluorescence with 473-nm laser light through all 12 fibers in parallel and analyzing the fluorescence time course in the 12 regions of interests (ROIs) on the camera images representing the fiber channels, we simultaneously recorded calcium signals in the entire set of brain regions. Mice were head-fixed, awake and engaged in a go/no-go texture discrimination task (see below). Across regions, we observed robust fluorescence changes (ΔF/F) with diverse time courses and signal amplitudes of up to 20% (**Fig. 1f**). Several control experiments confirmed that the observed signals represent calcium signals evoked by neuronal population spiking activity. First, excitation at 425 nm, a wavelength close to the isosbestic point of GCaMP6^31^, produced flat ΔF/F signal traces with significantly lower variance compared to 473-nm excitation (**Fig. 1g**), excluding major contributions of intrinsic signal changes or motion artifacts^26^. Second, we validated that fiber-optic signals reflect neuronal spiking by simultaneously recording single-unit activity with four tetrodes placed around a single fiber tip. Trial-related fiber-optic ΔF/F signals were well reconstructed by a weighted sum of the spike rates recorded from single units near the fiber tip (**Fig. 1j**; taking into account the GCaMP6 decay characteristics by convolution with an appropriate Kernel; see **Supplementary Notes** and **Supplementary Figure 2**). Third, we combined photometry with two-photon microscopy and found that brief electrical stimulation of superficial cortex elicited exponentially decaying calcium transients in GCaMP6m-labeled cortical neurons in the vicinity of the fiber tip and also resulted in a clear photometric calcium signal (**Supplementary Notes** and **Supplementary Figure 3**). Finally, we verified that signal cross talk between fiber channels is small, on the order of few percent for the closest fiber channels (**Supplementary Notes** and **Supplementary Figure 4**).

**Figure 2.**
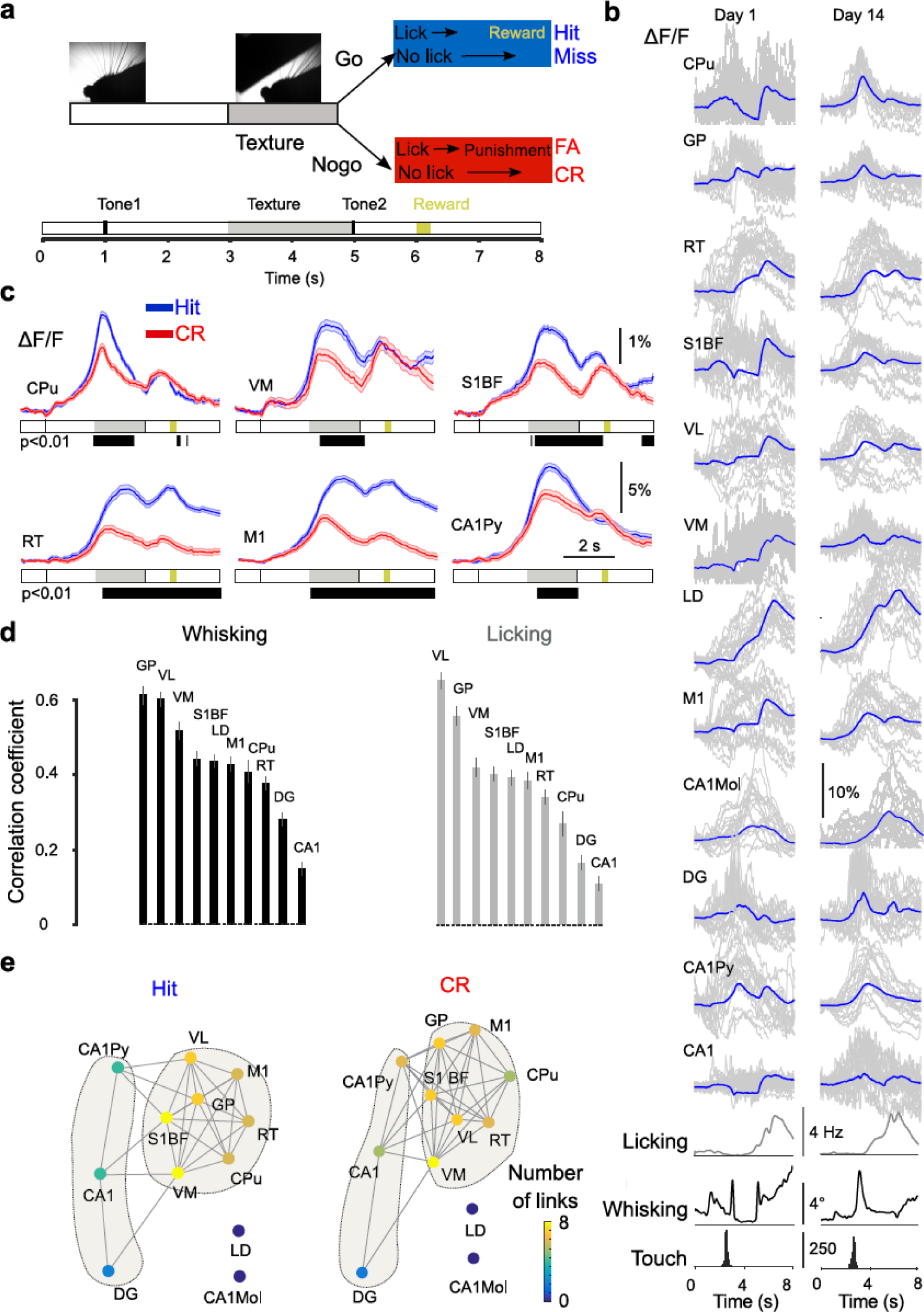
Large-scale network dynamics across multiple brain regions during texture discrimination behavior. (**a**) Schematic representation of the go/no-go texture discrimination task. (**b**) Example calcium signals recorded from 12 brain areas during Hit trials for day 1 and day 14 (blue curves are mean ΔF/F traces averaged over 303 and 315 Hit trials, respectively; example traces of individual trials are shown in grey; note that some individual traces are partially truncated). Traces of be-haviorally relevant parameters are shown below. (**c**) Example of trial-averaged ΔF/F signals for Hit (n = 315 trials) and CR (n = 323 trials) (mean ± SEM, mouse m7; grey bar below calcium traces marks period of texture presentation; black bars indicate periods of significant Hit/CR difference, p < 0.01; one-way ANOVA). (**d**) Correlation of calcium signals to whisking envelope and licking rate in texture discrimination trials (mean ± SEM, for n = 6 mice; Online Methods). (**e**) Functional network representation during Hit and CR trials using ‘force-directed placement’ for the session used to plot Figure 2c. Region nodes are coloured according to their number of links. Note that a higher density of links affects spacing between areas, i.e. multiple interlinked areas are pulled together into clusters.

**Figure 3.**
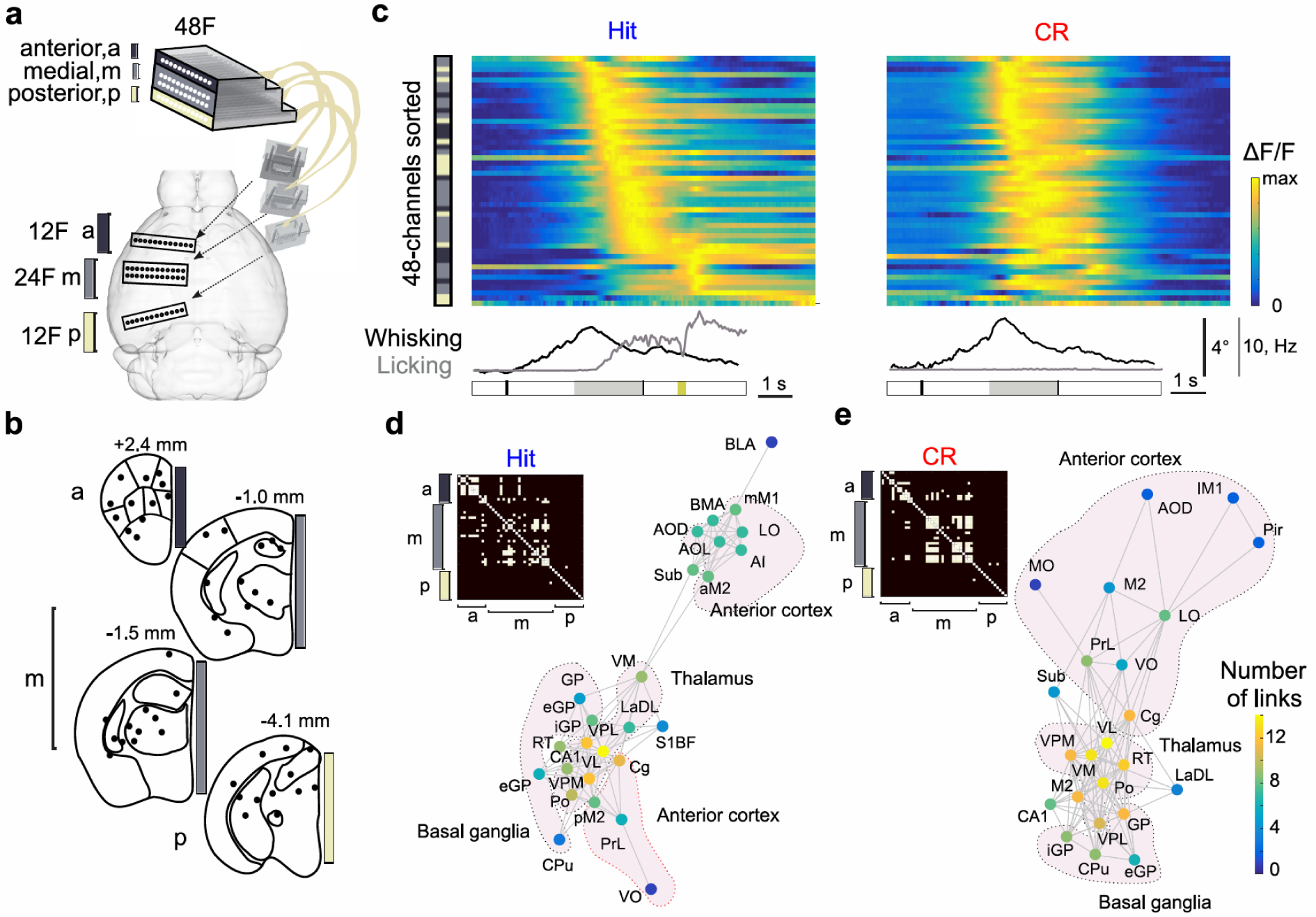
Simultaneous 48-channel multi-fiber photometry during texture discrimination. (**a**) Top view of a mouse brain surface and implanted multi-fiber arrays. (**b**) Schematic plot of the distribution of fiber tips across areas in anterior cortex (‘a’), medial areas of the basal-ganglia-thalamo-cortical loop (‘m’), and visual-hippocampal areas in the posterior section (‘p’). Numbers indicate rostrocaudal position relative to bregma. (**c**) Heatmap of trial-averaged R-CaMP1.07 signals for Hit (n = 244 trials) and CR (n = 210 trials). Average ΔF/F traces were normalized to their peak values. Traces are sorted according to the timing of their peak value. Left vector depicts the distribution of anterior, medial and posterior regions using same colour code as in (a). Bottom: mean whisking envelope and mean licking rate for Hit and CR trials. The horizontal bar plot shows a trial structure according to Figure 2a. (**d**) Top left: the adjacency matrix computed with pairwise correlation coefficients (only significant correlations, p < 0.05, were included in the plot). Functional network as in Figure 2e reconstructed for Hit trials. Areas (nodes) are coloured by the number of links. (**e**) Same plots as in (d) for CR trials.

### Multi-fiber photometry of behavior-related neuronal dynamics in 12 to 48 brain regions

Complex behaviors require coordinated activity in neuronal circuits spanning many brain areas. To exemplify how multi-fiber arrays can be applied to assess behavior-related large-scale circuit dynamics we recorded from multiple distributed brain regions in mice engaged in a go/no-go texture discrimination task^33^. The six mice implanted with 12-fiber arrays were habituated to head fixation and trained to lick for a water reward for one type of texture (Hit trials) and to refrain from licking for a non-rewarded different texture (correct rejection, CR, trials) (**Fig. 2a**; Online Methods). This sensorimotor task likely involves various subcortical regions, including basal ganglia nuclei as well as thalamic and hippocampal subregions, which all are known to interact with neocortical areas^34^. We therefore targeted the fiber tips to caudate putamen (CPu) and globus pallidum (GP), to several thalamic nuclei (lateral dorsal, LD; ventral lateral, VL; ventral medial, VM; and reticular nuclei, RT), to dorsal hippocampal regions (several sublaminae of CA1 and dentate gyrus, DG), and to neocortical regions (primary somatosensory barrel cortex, S1BF, and posterior part of primary motor cortex, M1) (a full list of regions targeted in this study and their abbreviations is provided in **Supplementary Table 1**).

In each mouse we recorded multi-regional calcium dynamics and revealed trial-related ΔF/F signals above baseline in all regions (**Fig. 2b**). Notably, it was possible to perform multi-fiber photometric measurements chronically over several weeks (2 - 6 weeks in 6 mice), confirming the stability of the implants. The time course of calcium signals was diverse and variable, typically showing prominent peaks in the early active touch phase (e.g. in CPu, GP, VL, DG, S1BF and M1) and/or in the later response phase, including licking action and reward retrieval (e.g. in LD, RT, and CA1). Calcium signals diverged for Hit and CR trials in all regions after touch onset, consistent with early discriminability^9^ (**Fig. 2c**). Furthermore, calcium signals in ventral thalamic nuclei (VM, VL) as well as GP were most strongly correlated to relevant behavioral variables such as whisking envelope and licking rate whereas hippocampal regions (DG and CA1) displayed only weak correlations (**Fig. 2d).**

High-density multi-fiber photo-metry allowed us to reconstruct a functional network of interacting brain regions. To estimate functional connectivity we used pairwise correlations between calcium signals^12^. A link was drawn between regions only for significant correlation coefficients (p < 0.05) and the resulting network was visualized with ‘force-directed placement’ (**Fig. 2e**; Online Methods). We calculated separate networks for Hit and CR trials based on the ΔF/F signals from the entire trial duration. We found that the thalamic nuclei (VM, VL) as well as S1BF took a central position in the network, possessing a higher number of links compared to other areas. This example illustrates how multi-region photometry can be applied to reveal functional networks recruited during a particular behavior.

To demonstrate that multi-fiber photometry can be scaled up for investigation of even larger brain networks, we in addition implanted one mouse with 48 fibers in one hemisphere. Three front pieces were implanted: one 24-fiber array at 1.4 mm posterior to bregma (including the same brain region targets as in the 12-fiber experiment), and two additional 12-fiber arrays at 2.4 mm anterior and 4.1 mm posterior to bregma, respectively (**Fig. 3a**). All fiber bundles were combined at the back end in a 48-fiber connector to couple all 48 channels to the photometry system. In this experiment, we used viral expression of the red calcium indicator R-CaMP1.07 instead of GCaMP6m. We trained this mouse in the texture discrimination task and recorded calcium signals in the mesoscale network, now including additional areas in the anterior cortex (mostly anterior motor cortex and prefrontal areas) and posterior regions (visual cortex, ventral hippocampus and superior colliculus; **Fig. 3b**). Trial-averaged ΔF/F calcium signals in many regions peaked during the texture presentation time (**Fig. 3c**; example Hit and CR trials for all regions are shown in **Supplementary Figure 5**). In addition, multiple cortical and subcortical areas displayed increased population responses upon reward delivery. These preliminary observations indicate that a highly distributed network is recruited in the texture discrimination task. As proof-of-principle this experiment demonstrates the power of high-density fiber arrays for scaling up fiber photometry to entire brain networks.

We again used pairwise correlations to quantify functional connectivity and to reconstruct 48-region functional networks for Hit and CR trials (**Fig. 3d**). Interestingly, these functional networks formed several distinct clusters (anterior cortical areas, thalamus, and basal ganglia) with features matching known anatomy. For example, one cluster contained mostly thalamic nuclei and was positioned between basal ganglia and anterior cortical areas, reflecting the anatomical organization of the basal-ganglia-thalamo-cortical loop^32^. Specifically, VL and VM were located on the shortest path between basal ganglia and anterior cortex, in line with anatomical evidence that VM/VL diffusely project to multiple anterior cortical regions^34^ in addition to sensory cortices. We conclude that multi-fiber photometry experiments enable identification and analysis of behavior-related functional networks across several tens of brain regions. Based on our network analysis one can predict that perturbing ventral thalamus (VM, VL) may have a strong effect on anterior cortical circuits. Such prediction can be tested by combining multi-fiber photometry with optogenetics through the same set of fibers.

### Combining multi-fiber photometry with optogenetic perturbation

Fiber-optic implants are widely used in optogenetic excitation or inhibition experiments^24,25^. To demonstrate that multi-fiber photometry can be combined with targeted optogenetic perturbations, we added a second laser beam path with a spatial light modulator (SLM) to our setup (**Fig. 4a**). The SLM can create multiple beamlets that can be coupled to a selected subset of fiber channels (**Fig. 4b**; Online Methods). This approach enables optogenetic perturbation of either individual regions or multiple regions together. Here we used VGAT-ChR2 EYFP transgenic mice, in which local neural circuit activity can be inhibited by blue-light excitation of ChR2-expressing GABAergic interneurons (Online Methods). In these mice, we virally expressed R-CaMP1.07 in the brain regions targeted by the multi-fiber array. Our goal was to perturb part of the functional brain network during texture discrimination behavior to induce changes in brain network dynamics and possibly behavior.

**Figure 4.**
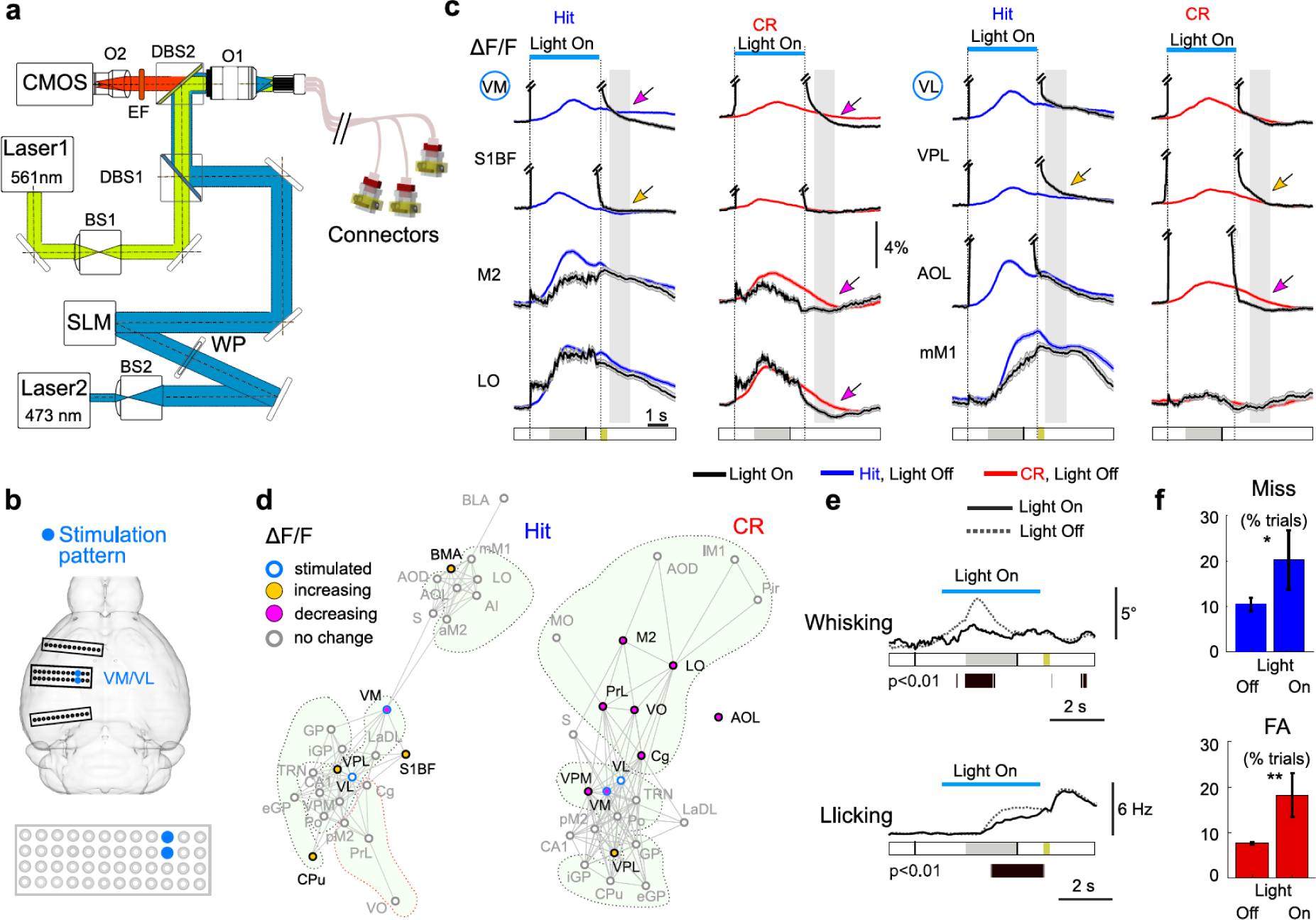
Simultaneous optogenetic perturbation of VL/VM thalamic nuclei combined with multi-fiber photometry. (**a**) Schematic of the optical setup adapted for optogenetic experiments. Laser 1 is used to excite R-CaMP1.07 fluorescence and laser 2 for ChR2 activation. Beam 2 is shaped using a standard 5x beam expander (BS2), a half-wave plate (WP) and a spatial light modulator (SLM) to create the required profile of beamlets in the objective (O1) plane for optogenetic perturbation. One dichroic beam splitter (DBS1) combines lasers for photometry and optogenetics. The second beamsplitter (DBS2) separates excitation and fluorescence light. (**b**) Position of fibers targeting VL and VM in the fiber implants on the skull. Schematic representation of the light pattern targeting both VL and VM through the 48-channel fiber array. (**c**) Trial-averaged ΔF/F traces for example brain regions (mean ± SEM, mouse m2). Traces for perturbed trials (‘Light On’) are plotted as solid black lines (n = 58 Hit trials and n = 44 CR trials, respectively). Unperturbed Hit trials are shown in blue (n = 244) and unperturbed CR trials in red (n = 210). Arrows highlight regions, in which calcium signals significantly diverged from unperturbed ΔF/F levels after perturbation. Grey vertical bars indicate the time window for post-perturbation analysis. (**d)** Schematic representation of the mesoscale network effect by VM/VL perturbation for Hit (left) and CR (right) trials. Functional networks shown in grey are taken from Figure 3d and e. Regions with reduced ΔF/F level in the late analysis window are shown in magenta; regions with increased ΔF/F are shown in yellow; regions with optogenetic perturbation in blue and unaffected regions as hollow grey circles. (**e)** Reduction in whisking envelope and licking rate upon VM/VL perturbation. (**f**) Increase in the rate of Miss and FA trials upon optogenetic perturbation in VM and/or VL. Miss rate (Miss) is plotted in blue and FA rate in red (* 0.01 < p < 0.05, ** p < 0.01; t-test; Online Methods).

We combined multi-fiber photometry and optogenetic perturbation in 3 mice: one mouse with the 48-fiber implant and two additional mice with 12-fiber array implants. With the SLM we shaped the laser beam wavefront such that the resulting pattern of beamlets targeted the thalamic nuclei VL and VM (either as a region pair or individually; **Fig. 4a,b**). We chose these regions as targets for perturbation because of the strong correlation of their activity with behavioral variables and their central position in the network (**Fig. 2e** and **Fig. 3d,e**). While mice performed the texture discrimination task we optogenetically excited GABAergic interneurons in VL/VM in random 30% of trials. We used a 4-s long illumination period starting before the first whisker-texture touch occurred and lasting until 1 s after texture withdrawal. Stimulation consisted of a 20-Hz train of 20-ms light pulses at a power density of ∼480 mW/mm^2^ at each fiber tip. Two issues restricted the temporal window, in which we could evaluate network effects of the optogenetic inhibition of VM/VL. First, the illumination power for optogenetic stimulation was ∼14-fold higher compared to the continuous 561-nm laser illumination used to excite R-CaMP1.07 fluorescence for the photometric recordings (∼1.3 mW/mm^2^ power density per fiber channel). As a consequence, the fiber channels targeted for optogenetic stimulation as well as neighboring fiber channels exhibited large signal increases during the stimulation period, presumably reflecting auto-fluorescence excited at 473 nm (**Fig 4c**; more distant fiber channels were less affected). Second, R-CaMP1.07 (as a R-GECO1 derivative) might be photo-activated by blue light^35^, which could lead to misinterpretations of fluorescence changes in terms of calcium dynamics. We checked this potential problem by applying brief 50-ms illumination pulses at 473 nm in a wildtype mouse expressing only R-CaMP1.07. This stimulation indeed elicited brief fluorescence transients which, however, decayed back to baseline rapidly (0.20 ± 0.08 s time constant, 9.1 ± 1.4% amplitude at 480 mW/mm^2^ power density; n = 100 pulses; n = 3 regions in one mouse; **Supplementary Figure 6**). To avoid signal contamination during light stimulation as well as transient photo-activation of R-CaMP1.07, we therefore restricted our analysis of ΔF/F signal changes to the time window from 0.5-1.5 s after the perturbation laser was turned off.

We evaluated the mean ΔF/F values in this analysis window for Hit and CR trials with and without perturbation of VM/VL (**Fig. 4c**; ‘Light On’ and ‘Light Off’ trials, respectively). For both Hit and CR trials we observed a significant ΔF/F reduction in VM and a trend towards reduction in VL. Several other regions were also affected. In particular, for CR trials and in line with our prediction (**Fig. 3d**) several anterior cortical areas reduced their activity after thalamic perturbation (**Fig. 4d**). Further signal changes occurred in S1BF and CA1 following individual perturbations of VM or VL (**Supplementary Figure 7**). Since all experimental parameters were the same in Hit and CR trials, the differential network effects suggest that optogenetic interference caused trial-type dependent changes in mesoscale circuit dynamics. While these results are preliminary, the main purpose of these experiments was to demonstrate the feasibility of combining multi-fiber photometry with optogenetics.

We also asked whether thalamic perturbations affect behavior. Indeed, we found significant decreases in whisking during the touch phase and in licking behavior (**Fig. 4e**; **Supplementary Fig. 7**), consistent with the strong correlation of VM/VL calcium signals with these variables (**Fig. 2d**). Finally, we examined the effect on task performance. All mice showed increased fractions of FA and Miss trials upon perturbation of the ventral thalamus (**Fig. 4f**; Online Methods). The higher fraction of FA trials and an increased licking rate during FA trials (**Supplementary Figure 7**) indicate that impairment in behavioral performance is not simply explained by suppressed motor function. We conclude that high-density fiber arrays enable optogenetic manipulations in individual or multiple brain regions. Combined with multi-fiber photometry, this approach makes it possible to first identify relevant regions in a mesoscale network and then to study how perturbation of these regions affects the functional networks and behaviour.

### Multi-fiber photometry in freely moving mice

The light-weight fiber bundles and miniaturized fiber-arrays also facilitate freely moving experiments. Here we demonstrate this opportunity with proof-of-concept measurements of mesoscale network dynamics in mice freely exploring a new environment and a novel object (Online Methods). We compared activity changes in different brain regions during distinct types of behavior (novel object approach and rearing) to baseline behavior (free cage exploration) (**Fig. 5a**). Upon object approach ΔF/F significantly increased in S1BF, CPu, GP, in basolateral/basomedial amygdala (BLA/BMA), and in ventral posterolateral/posteromedial thalamic nuclei (VPL/VPM) (**Fig. 5b**). Rearing, on the other hand, was correlated with increased ΔF/F values in DG and interestingly also with decreased ΔF/F values in S1BF and CPu as compared to baseline behavior (**Fig. 5c**). Obviously, further experiments and a more quantitative parametrization of behavioral variables will be required to relate mesoscale network dynamics to specific aspects of naturalistic behaviors^36^.

**Figure 5.**
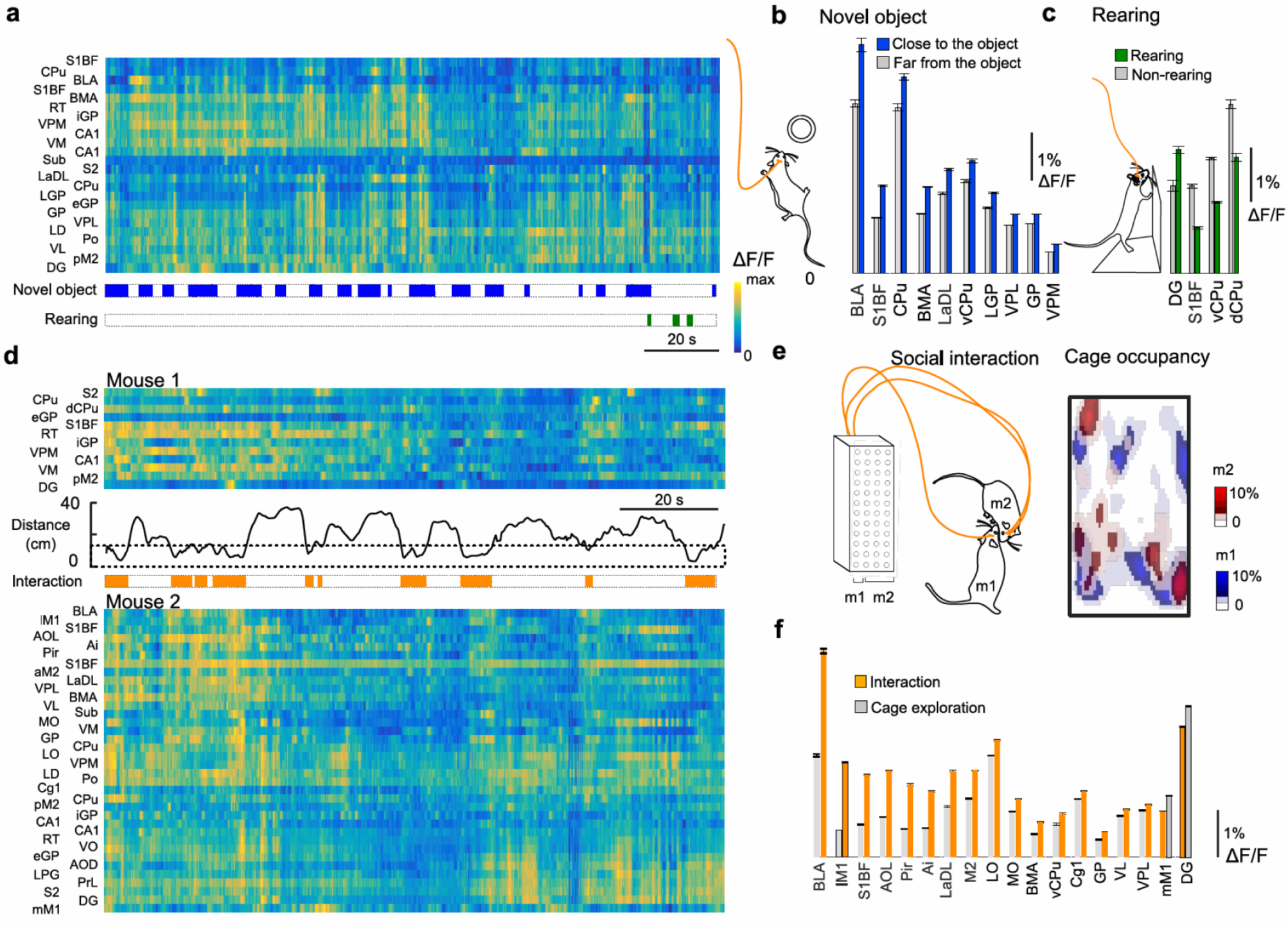
Multi-fiber photometry in freely moving mice. (**a**) Multi-fiber photometry in a mouse freely exploring the cage, sometimes approaching a novel object and sometimes rearing. Bottom: blue bars mark time periods of object approach (< 20 cm distance), green bars indicate rearing events. (**b**) Regions displaying significant changes in ΔF/F upon approaching the object compared to a time periods of free-cage exploration (p < 0.01; one-way ANOVA, mouse m2; additionally, the mean ΔF/F level had to be at least 0.5% higher than the median ΔF/F level during free cage exploration). (**c**) Regions displaying significant changes in ΔF/F for rearing events as compared to a time periods of free-cage exploration (p < 0.01; one-way ANOVA, mouse m2). (**d**) Simultaneous fluorescence recording in 48 channels from two socially interacting mice (m1: 12 fibers; m2: 36 fibers). (**e**) Left: schematic interaction of the pair of mice. Right: cage occupancy (m1 in blue and m2 in red). (**f**) Regions displaying significant changes in ΔF/F for time periods when mice were interacting (< 10 cm distance, ‘Interaction’) versus when they were far apart (> 20 cm, ‘Cage exploration’) (p < 0.01; one-way ANOVA test for n = 2 mice, anterior regions from mouse m2; additionally, the mean ΔF/F level during social interaction had to be at least 0.5% higher than the median ΔF/F level during free cage exploration).

We also performed multi-fiber photometry in two interacting mice by connecting separate branches of the fiber bundle to two animals and coupling the fluorescence signals to one photometry setup. Two male mice (one with a 12-fiber implant, the other with a 36-fiber implant) were placed in the same cage and approached each other multiple times during a session (**Fig. 5d)**. Several regions in amygdala were recruited during these social interactions (BLA, BMA and a dorsal part of lateral amygdaloid nucleus LaDL, p < 0.01; one-way ANOVA test for ΔF/F distribution of samples for a relative distance between two mice ‘far’, i.e. > 20 cm, or ‘close’, i.e. < 10 cm; **Fig. 5d, e**). Cortical regions (anterior lateral motor cortex (lM1), insular cortex (Ai), secondary motor cortex (M2), lateral and medial orbital cortices (LO and MO), S1BF and anterior olfactory nucleus (AOL), piriform cortex (Pir) and subcortical regions (CPu, GP; ventral thalamus VL and VPL) were recruited upon the close encounter with the other animal (significantly increased ΔF/F; p < 0.01, one-way ANOVA test; **Fig. 5f**). Consistent with recent studies^37^, it is likely that BMA and BLA neuronal ensembles encode context-specific information relevant for social interaction, while other areas such as S1BF and CPu may reflect multiple aspects of social touch and olfactory processing initiated by social interaction. Taken together, application of high-density fiber arrays in freely moving animals may facilitate our understanding of long-range circuits underlying complex naturalistic behaviors, including social interactions.

## DISCUSSION

We have presented high-density multifiber arrays as a novel type of neural interface that provides ample new opportunities for readout and optical control of large-scale mesoscale network dynamics in mammalian brains. We demonstrated simultaneous recording through up to 48 fiber channels from multiple subcortical and cortical regions in the mouse brain and revealed widely distributed network activity during texture discrimination behavior. We also established all-optical mesoscale network interrogation by optogenetically perturbing ventral thalamic nuclei, which we identified as central nodes in this behavioral task. Thalamic perturbations caused wide-spread changes in mesoscale network dynamics, especially in brain regions known to have anatomical connections with thalamic nuclei^32,34^, and also affected behavioral performance. Furthermore, we gave proof-of-concept that multi-fiber arrays can be applied in freely moving mice during natural behavior. These demonstrations highlight the utility and versatility of the multi-fiber arrays and we see great prospects for many further applications.

We believe that the number of fiber channels can be further increased. Because of their compactness and low weight, even implanting 6 front pieces into a mouse or rat brain should be feasible, particularly as they could be distributed across both hemispheres and possibly the cerebellum, too. If each front piece would carry 24 fibers, this approach could boost the number of simultaneously recorded brain regions into the range of 100-150. The fibers we used here were well tolerated by the tissue and we think that brain damage associated with multifiber implantations can be kept at a bearable level. For example, compared to a 3-mm cannula window implanted above CA1—a widely applied method for hippocampal imaging that requires tissue aspiration^38^—the total volume of inserted fiber material in our 48-channel experiment was about 5-fold smaller (∼1.2 mm^3^ compared to ∼7.7 mm^3^). To make fiber probes even less invasive thinner optical fibers or tapered fiber tips^39^ could be used. To maximize the number of simultaneously imaged regions, it should also be possible to integrate multi-fiber arrays into wide-field microscopes^12,14–16^, implanting fiber arrays at an angle and using two cameras or dedicated segments of the same CMOS sensor for parallel multi-fiber and wide-field measurements. Many further opportunities exist for hybrid approaches, combining multi-fiber arrays with other existing technologies. For example, high-density fiber arrays could expand the use of fiber optics in fMRI experiments^40–42^ and thereby contribute to a better understanding of the relationship between neuronal, glial, and vascular dynamics. A combination with extracellular silicon probes^6^ or emerging optoelectronic neural probes^43,44^ may provide new possibilities for relating individual neuron activity to larger-scale population activity^45^, even up to the mesoscale^46^. Similarly, the combination with multi-site neurochemical recordings^47^ would allow studies of how neuromodulator activity impacts brainwide neural network dynamics.

Because of their simplicity multifiber arrays are especially well suited for applications in behaving animals. Here we have demonstrated applications in both head-fixed and freely moving mice. Multifiber photometry should be equally well applicable in rats, non-human primates^48^, and other mammalian species. Similar to single-fiber approaches, the multi-fiber approach will gain its full strength through the clever combination with modern genetic tools that permit cell-type or pathway-specific labeling with optical indicators, not only for neuronal calcium signals but also for membrane potential^21^, dopamine release^49^, glial dynamics^41,42^, metabolic substrates^50^, etc.. Likewise, multi-fiber implants could be useful in photo-activation experiments employing calcium integrators for marking of, or inducing targeted gene expression in, neuronal populations that are activated during specific time windows^51,52^. Multicolor approaches should allow the parallel usage of multiple indicator systems. Finally, as we demonstrated here in VGAT-ChR2 mice, the multi-fiber arrays can also be applied for multisite optogenetic control, benefitting from the large available palette of opsins^53^ and photoactivatable molecules. Many other possible combinations of activity reporters with optogenetic actuators exist. Suitable expression of the respective proteins throughout the brain may be achieved for instance in transgenic mouse lines^13^ with supplementary viral injections. To avoid the work load of multiple virus injections and to facilitate more complex expression patterns, it will be interesting to test virus release from silk fibroin drops^54^ applied to the fiber tips. The integration of optogenetic manipulations with multifiber photometry or wide-field imaging^12^ has the potential to enable all-optical studies on the mesoscale, complementing current efforts to establish all-optical interrogation at the microcircuit level with single-cell precision^55–57^.

We anticipate that high-density fiber arrays will become important tools to study the complex functional organization of the mammalian forebrain. Not only do they allow to reconstruct functional networks related to specific behavioural aspects, they also can be applied chronically to study functional network changes in a longitudinal approach, e.g., during learning or disease progression. Here we assessed functional networks using pairwise correlations between calcium signals as simple connectivity measure. Clearly, many further computational approaches such as Granger causality^58^ and information theory^59^ are available, which should be directly applicable to multi-fiber photometry data. State-of-the-art computational methods and graph analysis^60^ may help to identify characteristic activity patterns in the brain-wide circuits. This, in turn, will facilitate unbiased functional mapping of the brain networks in order to identify circuit components relevant under various healthy and diseased conditions.

## ACKNOWLEDGEMENTS

We would like to thank Hansjörg Kasper, Martin Wieckhorst and Stefan Giger for technical assistance, Jose Luis Alatorre Warren for the registration of histology sections to the Allen Brain Atlas and fiber shaft tracking, Alexander Jovalekic for help with electrophysiological recordings and spike sorting, Asli Ayaz, Ladan Egolf and Christopher Lewis for comments on the manuscript. This work was supported by grants to F.H. from the Swiss National Scien**c**e Foundation (310030B_170269) and the European Research Council (ERC Advanced Grant BRAINCOMPATH, project 670757), by a Transfer Project and an IPhD Project from SystemsX.ch (51TP-0_145729 and 51PHP0_157359, respectively), and through a Roche Joint Collaborative Project.

## AUTHOR CONTRIBUTIONS

Y.S. and F.H. designed the experiments, Y.S. conducted the experiments and analyzed data, M.C. assisted in fiber-optic experiments, L.S. performed brain histology, Y.S. and F.H. wrote the paper.

## METHODS

### Design of high-density multi-fiber arrays

We utilized components of MT fiber connector technology, specifically ferrules that can accommodate various numbers of fibers (8, 12, 24, and 48 fibers; part numbers 17185, 7413, 12599; US Conec, Hickory, NC). Optical fibers with polyimide coating (100-µm core diameter, 124-µm outer diameter, NA 0.22; UM 22-100, Thorlabs) were fitted into the ferrule. To mount all fibers such that they would target a specific set of brain regions, we aligned them to a custom-built template defining the required implantation depths. After alignment, we glued the fibers inside the ferrule with a two-component epoxy (302-3M 1LB Kit Part A and B; Epoxy Technology). An additional drop of epoxy was applied to the external surface of the array to be optically connected to the fiber bundle. We let the epoxy cure over 24 hours and then polished the connecting surface in consecutive steps starting with a rough diamond lapping sheet of 30-µm grit, to a silicon carbide lapping sheet of 5-µm grit, then gradually moving to a calcined alumina lapping sheet of 0.3-µm grit (from LF30D, LF5P, LF3P, LF1P to LF03P; Thorlabs). A custom-designed bare ferrule polishing puck was used to hold the ferrule orthogonal to the polishing pad (NRS913A, Thorlabs). Ethanol was added on to the polishing pad to reduce friction. After the polishing procedure we added alignment pins to the implant (part numbers 16741, 16742; US Conec). A custom-designed magnetic housing was added after the multi-fiber array was fixed to the skull with dental cement. We achieved a coupling efficiency of 75.2 ± 1.2% between a 12-fiber implant and the connecting bundle.

### Design of fiber bundles connecting to multi-fiber implants

The design of the fiber bundles connecting the multi-fiber implants to the fiber photometry setup followed similar steps regarding the fitting of fibers into the required ferrules. First, optical fibers (UM 22-100, 0.22 NA, 100µm core; Thorlabs) were cut to the required length with a ruby fiber scribe (S90R; Thorlabs). For the head-fixed behavior in the texture discrimination setup we used 2-m long fibers. For freely-moving experiments we prepared a 3-m long fiber bundle. To protect fibers we fitted 4 fibers each into a protective light-weight tubing (FT900Y; Thorlabs). After all fibers were positioned, we applied epoxy to the ferrules. To stabilize the fiber facet for subsequent polishing and ensure high optical quality for two connecting the two surfaces (multi-fiber implant and fiber bundle), we added an extra drop of epoxy on the surface connecting to the front piece. We polished both the proximal and distal end of the fiber bundle as discussed above. The proximal fiber connector was mounted in a custom-designed holder fitting in a 30-mm cage XY-translator for 1” diameter optics (CXY1; Thorlabs) to position it in the optical setup.

### Optical setup

The proximal end of the fiber bundle was integrated into an optical setup for calcium indicator fluorescence measurements and optogenetic manipulations. We used an Omicron LuxX 473-nm laser (or alternatively a Coherent OBIS LX 488-nm laser) for excitation of virally expressed GCaMP6m and activation of ChR2, and a Coherent OBIS LS 561-nm laser to excite R-CaMP1.07. To achieve stable CW operation, lasers were run at 80% of maximal output power. A variable neutral density filter (NDC-25C-4M; Thorlabs) was used to reduce fluorescence excitation power to ∼1.3 mW/mm^2^ per fiber channel at the fiber tip. One or two cylindrical lenses shaped the excitation beam in an appropriate illumination pattern at the object plane of the objective. First, we expanded and collimated the circular beam with an achromatic Galilean beam expander (GBE05-A; Thorlabs). Second, to create a line illumination pattern matching the 12-fiber array we used a 75-mm focal length cylindrical lens (LJ1703RM-A; Thorlabs) placed at ∼145 mm distance from the objective (TL4x-SAP; Thorlabs). For the 48-fiber array a rectangular illumination pattern was required. For this purpose we added a cylindrical lens with 150-mm focal length (LJ1629RM-A; Thorlabs) oriented 90° with respect to the 75-mm focal length cylindrical lens. A first dichroic beamsplitter (F38-495, AHF) combined the two laser beams along the optical axis. A second dichroic beamsplitter (F58-486 dual line, AHF) coupled the excitation light (473 nm and/or 561 nm) into the objective (TL4x-SAP, Thorlabs) and transmitted the fluorescence in the emission spectral windows for both GCaMP6m and R-CaMP 1.07. To separate fluorescence signals from residuals of the excitation light and to minimize auto-fluorescence generated in a broader spectral range we used emission filters (525/50 nm, F37-516, AHF, for GCaMP6m and 605/70 nm, F47-605, AHF, for R-CaMP1.07) and a triple line notch filter at 425, 473 and 561 nm (ZET405/473/561 for Omicron LuxX 473nm and Coherent OBIS LS 561nm lasers, or, alternatively, for a combination of ZET405/488 ZET488/561 Chroma Technology Corp. for Coherent OBIS LX 488nm and Coherent OBIS LS 561nm lasers). To image the fiber array end face onto the camera we used a 200-mm focal tube lens (Proximity Series InfiniTube) with internal focusing. The image was created at the back focal plane of the tube lens on the CMOS sensor (ORCA Flash4.0, Hamamatsu camera).

### Animals and surgical procedures

All procedures of animal experiments were carried out according to the guidelines of the Veterinary Office of Switzerland and approved by the Cantonal Veterinary Office in Zurich. A total of 22 male mice (2-6 months age) were used in this study. GCaMP6m expression was induced by AAV2.9-hSyn-GCaMP6m injections into the brains of C57BL/6 mice (n = 12) followed by implantation of a 12-fiber array. Mice were anesthetized with 2% isoflurane (mixed in pure oxygen). Body temperature was maintained at 37°C. To prevent inflammation and pain during anesthesia, a 0.1µl/g bw of Metacam was injected subcutaneously. Prior to implantation, connective tissue was removed from the skull bone. The bone was additionally polished, dried and iBond (Kulzer, Total Etch) was applied to ensure best adhesion of the skull to the connective dental cement. To further stabilize the implant on the skull we used Charisma (Kulzer, A1) to produce a thin ring on the skull rim. Both iBond and Charisma require UV light curing. Small slit-like craniotomies were made to allow for virus injections and implantation of fiber arrays (carried out on the same day). First, ∼120 nl of AAV2.9-hSyn-GCaMP6m were delivered into all 12 areas of interest at a rate of 20 nl/min. A custom setup with a syringe and a barometer was used to control pressure injections. In order to allow for local diffusion and to avoid possible refluxes, the glass injection pipettes (10-15 µm diameter) were kept in place after injection for 10 min. Second, a 12-fiber array was implanted +0.4 mm from midline tilted at an angle of 15 degrees. We oriented the fiber array such that the most lateral fiber efficiently targeted CPu (−1.06 mm from bregma) and the most medial fiber targeted hippocampal areas (CA1, DG; −1.46 mm). Prior to fiber implantation we slightly scratched the dura surface. After, the craniotomy was sealed with Vaseline, which melts at body temperature and completely covers the craniotomy. Next, we applied dental cement (Tetric EvoFlow A1) on the skull and around the implant followed by UV light curing. A light-weight metal head post was additionally cemented to the skull, allowing for painless head-fixation during the behavioral experiments. After two weeks of recovery, the mice were habituated to head-fixation and trained in the texture discrimination task.

Transgenic VGAT-ChR2-EYFP mice (n = 5)^62^ were injected with AAV2.1-EF1a-R-CaMP1.07 and implanted with multi-fiber arrays following the protocol described above. Four mice were implanted with single 12-fiber arrays. One mouse was implanted with 3 arrays: two arrays with 12-fibers each and one with 24-fibers (total of 48 channels). The single 12-fiber arrays were implanted in frontal and posterior regions (+2.4 mm and −4.1 mm from bregma, respectively) while the 24-fiber array targeted the same channels as in the 12-fiber array experiments (implanted at −1.46 mm from bregma) with additional fibers implanted in ventral striatum (vCPu), lateral globus pallidus (LGP), lateral thalamus (Po and VPL) and amygdala nuclei (BMA, BLA and LaDL).

To combine fiber photometry and extracellular electrophysiology two C57BL/6 mice were implanted with an opto-tetrode using a similar preparation protocol as for the multi-fiber array. Mice were injected in posterior M1 with 120 nl of AAV2.9-hSyn-GCaMP6m. The opto-tetrode was implanted at around 350 µm depth in posterior M1 and a ground screw was placed contralateral to the implant hemisphere (see **Supplementary Notes** for further details). To estimate the level of immune-response induced by the multi-fiber array, we implanted one transgenic CX3CR1–EGFP mouse with a multi-fiber array containing four UM22-100 optical fibers. Similarly, one C57BL/6 mouse was used for simultaneous histological staining of astrocytes with GFAP and of microglia with Iba1 (see below and **Supplementary Figure 1)**. To estimate the level of R-CaMP1.07 fluorescence (and test for potential photo-activation) with 473-nm excitation light (as used for optogenetic perturbation) we injected AAV2.1-EF1a-R-CaMP1.07 in one C57BL/6 mouse into the same set of medial regions as in 12-fiber experiment.

### Texture discrimination task

Following fiber array implantation and two weeks of recovery, mice were first accommodated to head fixation through a series of short-duration head fixations. After starting water scheduling, each mouse were first trained to lick upon a texture presentation. After this shaping period we added presentations of either the go-texture (sandpaper grit size P100) or the no-go texture (sandpaper grit size P1200) and trained the mouse to discriminate the two texture types^33^. For the go-texture the mouse had to lick at a water spout to receive a water droplet as reward (Hit trial). Failure to lick was considered a Miss trial. For the nogo-texture the mouse had to refrain from licking so that lick/no-lick events were interpreted as false alarm (FA) and correct reject (CR) responses, respectively. The two texture types were presented with 50% probability. Each trial started with a TTL pulse to synchronize the calcium recording with the behavioral setup. One second after trial start, a first 2-kHz auditory tone (2 pulses of 100-ms duration at 50-ms interval) signalled the start of the texture approach (∼90° to the whisker pad). It took 2 s for the texture to reach the end position so that the first whisker-to-texture touch occurred in the interval 2–3 s after trial start. Textures stayed in contact with the whiskers for 2 seconds and were retracted afterwards. Upon texture retraction a second 4-kHz auditory tone (4 pulses of 50 ms duration at 25-ms interval) signalled the start of 2-s report period, during which a water drop was delivered if the mouse licked correctly for the go-texture (Hit trial). A loud white noise sound stimulus of 4-s duration was presented as punishment if the mouse licked for the nogo-texture (FA trial). Both reward and punishment were omitted if the mouse did not lick (CR and Miss trials for nogo-texture and go-texture presentations, respectively). The lick detector was reachable throughout the entire session. Textures were presented pseudo-randomly with no more than 3 presentations of the same texture type in 3 consecutive trials. After learning the texture discrimination task mice reported their decision by starting to lick during the late period of a texture presentation (3.5-4 s after trial start) and then during the response period (5-7 s after trial start).

### Optogenetic manipulation of mesoscale circuit

To perturb dynamics in the mesoscale circuit we used VGAT-ChR2 EYFP transgenic mice. A 473-nm laser (Omicron LuxX) was used to excite GABAergic neurons. The beam was expanded (5x GBE05-A, Thorlabs) and directed onto a spatial light modulator (SLM SN 4479, Meadowlark Optics) to shape the wave-front of the laser beam for patterned illumination. To achieve a pure voltage-dependent phase shift, the linear polarization of the incident laser beam should be aligned parallel to the extraordinary axis of the SLM pixels, which we achieved with a zero-order half-wave plate (WPH10M-473, Thorlabs). A programmable phase shift pattern can then be created by changing the voltage on the corresponding pixels. The phase shift pattern across the SLM was optimized with the Gerchberg-Saxton algorithm to generate multiple beamlets in the object plane of the objective for coupling the excitation light into the fiber channels selected for brain region perturbation. We applied a 4-s long period of 473-nm stimulation from 2-6 s after trial start using 20-ms pulses at 20 Hz produced by a waveform generator (Agilent 33500B; TTL-triggered from the DAQ board for behavior control).

### Post-hoc immunohistochemistry

One to two months after training, mice were anaesthetized (100 mg /kg bw ketamine and 20 mg /kg bw xylazine) and perfused transcardially with 4% paraformaldehyde in phosphate buffer, pH 7.4. After perfusion, tissue was removed from the skull and the head including the multi-fiber implant was additionally fixated in 4% paraformaldehyde for one week. Then, the ventral (bottom) side of the skull bone was removed and the brain was carefully extruded. Coronal sections (75 – 100 µm thickness) were cut with a vibratome (VT100, Leica). Prior to mounting, coronal sections were blocked in 10% normal goat serum (NGS) treated with 1% Triton at room temperature and incubated overnight at 4°C in 5% NGS, 0.1% Triton, and antibodies against GFAP (mouse monoclonal antibody; 1:500, Sigma 032M4779) and Iba1 (rabbit polyclonal antibody; 1:500, Wako Chemicals 019-19471) for astrocyte and microglia staining, respectively. Stained sections were mounted onto glass slides and confocal images were acquired with Olympus FV1000.

### Registration to the Allen brain atlas and tracking of optical fibers

Image processing was performed on grayscale confocal images of the coronal histological sections. To obtain a sharp edge at the brain’s midline in each coronal slice, all images were manually segmented from the background using the *Lasso* segmentation tool of the software Avizo 9.3.0 (FEI, Hillsboro, Oregon). The pre-processed images were then registered in elastix^63^ software using the following steps. One of the central sections that contained hippocampus was used as a first reference. The images of two directly adjacent sections (anterior and posterior) were registered to the central image using a 2D rigid-body transformation and the mean squared difference as similarity measure. The images aligned such were then used as reference themselves and their adjacent sections registered. This procedure was repeated until all sections were aligned. Using MATLAB’s DIPimage (www.diplib.org/dipimage) and tools for NIfTI and ANALYZE image^64^, a 3D NIfTI image was created for each mouse brain with inter-slice spacing between 0.075 and 0.1 mm. Each 3D NIfTI dataset was then registered to the Allen Mouse Brain volumetric atlas^32^ (downloaded from the Scalable Brain Atlas^65^) with the help of Avizo Software. The sharp midline edges of the brains and the atlas^66^ reference data imposed an additional anatomical constrain and thus improved the alignment. The registration was implemented in two steps, first using only rigid-body transformations (translation and rotation), then allowing for isotropic scaling and, if needed, additional anisotropic scaling. All registered slices were validated by a human. Mouse brain-to-atlas registrations were carried out by using Normalized Mutual Information^67^ as similarity measure and did not allow for shearing transformations. Once the 3D datasets were registered to the reference atlas space, the fibers were manually segmented in each slice using Avizo’s *Lasso* segmentation tool. 3D reconstructions of these segmentations allowed us to identify the fiber ends and obtain their coordinates in the atlas space.

### Network reconstruction

Adjacency matrices were constructed from pairwise correlation coefficients^68^. Cross-correlation values were normalized such that the autocorrelations at a zero time lag equal to one (*xcorr* function in MATLAB). The maximum correlation coefficient within a time lag window of ±250 ms was selected for analysis. To test for significant correlations we estimated p-values using t-statistics (*corrcoef* function in MATLAB). Significant correlations (p < 0.05) were chosen to construct the functional network. To visualize the resulting functional network on a 2D plane, we used a force directed placement algorithm^69^, which strives for uniform edge lengths and makes nodes that are not connected by an edge tend to be drawn further apart.

### Tracking freely moving mice

We tracked freely moving mice using custom-written MATLAB code. Video frames were read into a pre-allocated data array, which was reshaped according to the coordinates of the arena (specified polygonal region of interest). The image intensity values between minimum and maximum were normalized to the [0,1] range and binarized according to a 10% threshold. To account for few noisy pixels in the data, we removed small objects with areas containing less than 550 pixels from the binarized array. Area and centroid coordinates were measured for every frame with the *regionprops* function. In order to track two mice, centroid coordinates were assigned to the respective mouse according to the nearest centroid in the previous frame. The Euclidean distance to the object and to another mouse was calculated from the respective centroid coordinates. To create occupancy plots we calculated histogram bin counts of total 50 bins (5 x 10) and applied 2D Gauss-filtering (σ = 2 bins). Peaks in the histogram reflect dwelling spots in the arena and in the case of two mice also indicate places of interaction. Rearing behavior was difficult to identify with our code, we therefore manually found time stamps for rearing events.

## SUPPLEMENTARY INFORMATION

### Supplementary Notes

### Reconstruction of fiber-optic calcium signals from a local network of spiking neurons

To validate if fiber-optic calcium signals represent local neuronal population spiking activity, we combined fiber photometry with direct electrophysiological recordings using custom-designed opto-tetrodes (n = 2 mice). Four tetrodes were positioned and glued symmetrically around an optical fiber UM22-100 (**Supplementary Figure 2a**). This opto-tetrode was mounted on a micro-drive (AXONA A3324-94 HS132HN) such that it could be gradually lowered into the brain. Mice were injected locally into primary motor cortex with 120 nl of AAV2.9-hSyn-GCaMP6m and the opto-tetrode, consisting of the fiber tip and the attached 16 electrodes (Platinum 10% Iridium, California Fine Wire Company), was implanted in deep layers of posterior motor cortex. A ground screw was placed on the contralateral hemisphere. During measurements the implant was connected via the optical fiber to the optical setup designed to measure GCaMP6m fluorescence. In parallel, multi-unit activity was measured with the 16-channel AXONA pre-amplifier and the AXONA DAQ USB system.

The chronic implant allowed us to record from both the tetrodes and the fiber channel over three consecutive weeks. Mice were trained in the texture discrimination task. When a mouse reached expert level (i.e., above 80% success rate of Hit and CR trials), the implant was lowered in steps of 120 µm into the brain after each recording session. TINT Cluster Cutting and analysis software from AXONA were used to cluster spike data recorded on each tetrode. We used k-means algorithm to identify 4 to 5 clusters on every tetrode. Every cluster was manually validated by the shape of the spike waveform. We identified spurious spikes (spikes with onset times different compared to the average waveform) by setting an additional voltage threshold at early time points and clustering data along this dimension. Spurious spikes were clearly separable along this dimension and were manually excluded from the original cluster.

In total we collected recordings from 364 single units with simultaneous fiber-optic calcium recording (from 18 sessions in n = 2 mice, 62 single units from the first mouse and 302 single units from the second). The spiking rates of three example units during texture discrimination are shown in **Supplementary Figure 2b**). Some units were active during Hit trials (units 1 and 2) while others increased their spike rate in CR trials (e.g., unit 3). We collected single units across all behaviour sessions and merged them into one dataset, which was then used to reconstruct the fiber-optic calcium signal (**Supplementary Figure 2c,d**). We presume that this dataset was representative of spiking neurons dynamics in deep layers (M1) during the texture discrimination task. This dataset consisting of single units from each mouse was least-squares fitted to the calcium signal (for Hit and CR trials) recorded every trial. A weighted sum of single-unit spike rates, convolved with an exponentially decaying kernel function (500 ms time constant, approximating the calcium indicator decay time course), described well trial-specific population dynamics (see cumulative R^2^ value for single units sorted by their contribution to the population signal; **Supplementary Figure 2d**). Interestingly, to achieve 90% of explained variance in a population calcium signal, more single units were required for Hit trials as compared to CR trials (24.2 ± 4.6 and 8.5 ± 2.4 respectively; mean ± standard deviation; **Supplementary Fig 2e**). Thus, the population calcium signal can be explained as a sum of the heterogeneous spiking dynamics in the local neural circuit. Although we cannot exclude the possibility that labelled neuropil also contributed to the fiber-optic calcium signal, there was no evidence of an extra signal component that could not have been explained with the spike rates in the behaviourally relevant single units.

### Comparison of fiber photometry and two-photon calcium imaging

In order to further verify our interpretation of the photometry signal as population average of cellular calcium signals we also performed fiber photometry under a two-photon microscope. The optical fiber (UM 22-100) and a piggyback stimulation electrode (∼10 µm diameter glass pipette filled with PBS; extending ∼100 µm from the optical fiber tip) were inserted into the neocortex underneath the objective of a two-photon microscope system (Ti:sapphire laser system, Mai Tai HP, Newport Spectra Physics; Nikon LWD 16x/NA0.8 objective; galvanometric scan mirrors model 6210 Cambridge Technology; Pockel’s cell, Conoptics). The optical fiber and the electrode were glued together and attached to a micromanipulator (Luigs and Neumann Mini 25-XL). In acute experiments, mice were anesthetized with 1.5% of isoflurane. The fiber-electrode combination was inserted into somatosensory cortex through a mm-sized craniotomy (**Supplementary Figure 3a)**. We imaged neurons expressing GCaMP6m calcium indicator with the two-photon microscope and, simultaneously, recorded fluorescence signals via the UM22-100 fiber connected either to our photometry setup or to a simple detection module (GFP filter F37-516, AHF; and a PMT H7422-40, Hamamatsu). We activated a local population of neurons by injecting a 20-µA, 50-ms current pulse through the stimulation electrode. Using this approach we were able to compare the ΔF/F signals and the spatial distribution of activated neurons as measured by the two-photon microscope objective and as seen by the optical fiber: Injection of a current into superficial somatosensory cortex evoked a responses in 5 to 10 cell bodies and throughout the local neuropil approximately 70 µm away from the fiber tip (**Supplementary Figure 3b**). We estimate that the fluorescence collection volume of the fiber channel to cover approximately 120 µm^3^ (roughly defined by the fiber diameter). When the optical fiber was used to collect two-photon excited fluorescence, we could detect only photons from within the volume defined by the fiber diameter and its numerical aperture (**Supplementary Fig 3c**; note the example neuron outlined with a dashed line detected by two-photon imaging but invisible to the fiber). The time course of calcium signals upon two-photon excitation (detected either in the two-photon microscope or through the optical fiber) was also comparable to the time course of the calcium signals measured with one-photon excitation and fiber photometry (**Supplementary Figure 3d**). Calcium signal amplitude also depended on stimulation intensity (**Supplementary Figure 3e**). We conclude that the fiber-optic calcium signals nicely reflect the population averaged calcium transients observed through a two-photon microscope.

### Evaluation of signal cross talk between adjacent fiber channels

How far can fluorescence light generated at one fiber tip still be seen by neighboring fibers? To estimate cross-talk between fiber channels we sequentially illuminated single fiber channels while measuring fluorescence signals in all channels (**Supplementary Figure 4a**). We normalized fluorescence signals (or their variance) generated by neighboring fibers to the fluorescence generated by direct illumination and analyzed their distance-dependence (**Supplementary Figure 4b**). In the closest fiber tips (0.5-1 mm distance) we found about 5-10% fluorescence contribution of the neighboring fiber channel whereas values rapidly declined to negligible levels with distance (**Supplementary Figure 4c**). Two effects may contribute to the cross talk: i) Propagation of the excitation light from the excited channel to neighboring fibers, generating fluorescence underneath their tips; ii) Propagation of fluorescence light excited at the tip of excited fiber to neighboring channels. We believe that most of the bleed-through signal originates from cross-talk of excitation light, meaning that it actually represents additional fluorescence that is generated in the neuronal population under a fiber tip by diffuse excitation light from a neighboring fiber. This cross-talk of excitation light actually is less a problem because it does not confound the signal of interest and simply adds to the local signal intended to be measured. Spread of fluorescence light generated under a neighboring fibers presumably contributes less, given the lower intensity of the fluorescence light compared to the excitation light. Our assessment thus indicates that the cross talk between fiber channels is small and that the high density fiber arrays enable multi-fiber photometry from selected subsets of brain regions with good regional specificity.

## SUPPLEMENTARY FIGURES

**Supplementary Figure 1.**
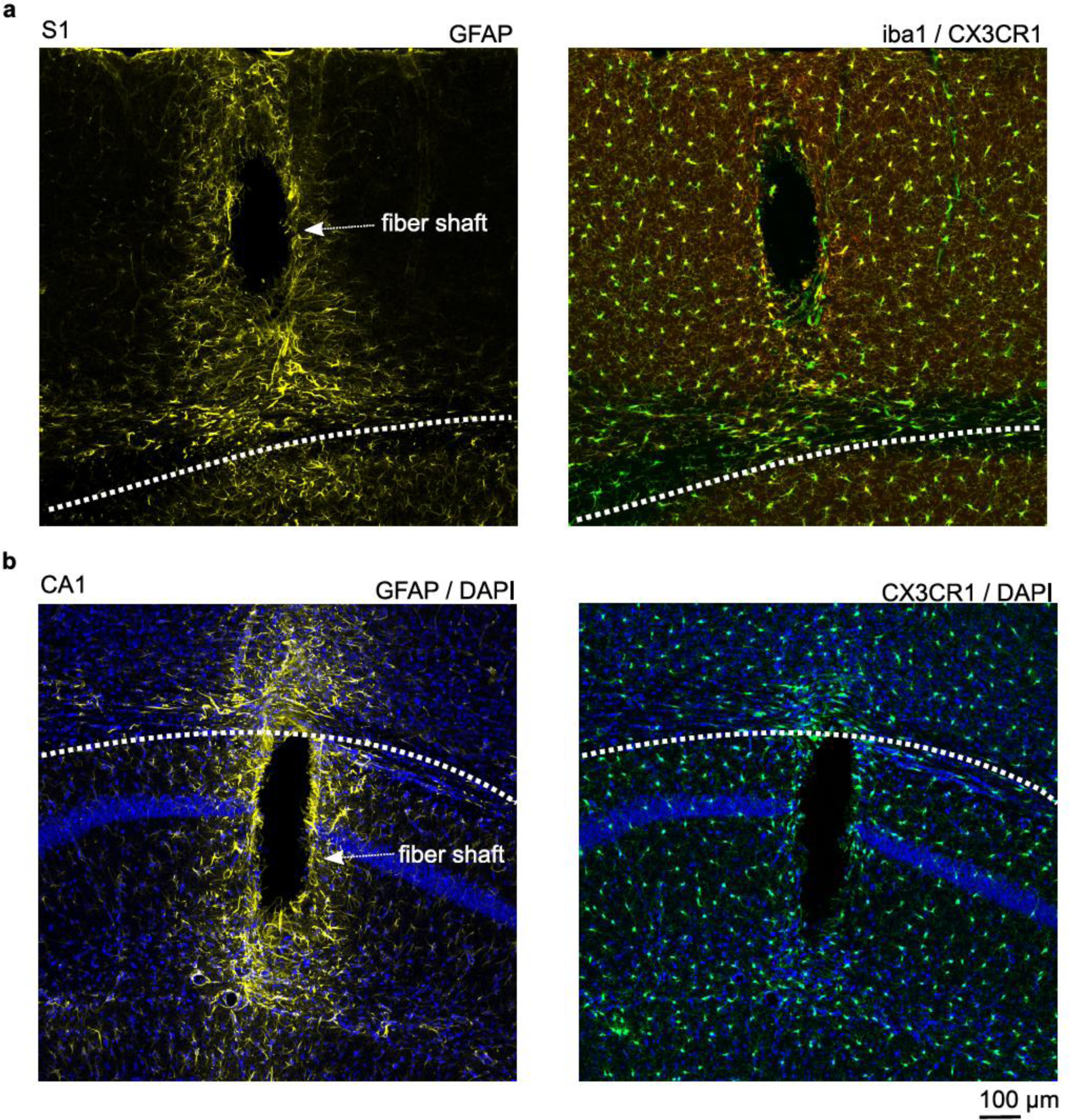
Glial response in mouse brain tissue 1 month after implantation of a UM22-100 multi-fiber array. (**a**) Left: Confocal image of astrocytes stained for GFAP (yellow) surrounding an optical fiber shaft in S1 cortex. A moderate astroglial scar formation is detected near the optical fiber. Right: Confocal image of microglia in S1 cortex (EGFP-expressing in a CX3CR1 mouse, green, and co-stained against Iba1, yellow). Note that microglia density is not markedly changed close to the fiber shaft as compared to more distant tissue areas. (**b**) Left: Confocal image of astrocytes surrounding the fiber shaft in the hippocampal CA1 region (yellow). The CA1 pyramidal layer is marked by dense band of DAPI-stained nuclei (blue). Right: Similar image with EGFP-stained microglia in the CX3CR1-EGFP mouse line. A moderate astroglial scar formation is detected near the optical fiber. Microglia density is not markedly changed, except for corpus callosum above CA1.

**Supplementary Figure 2.**
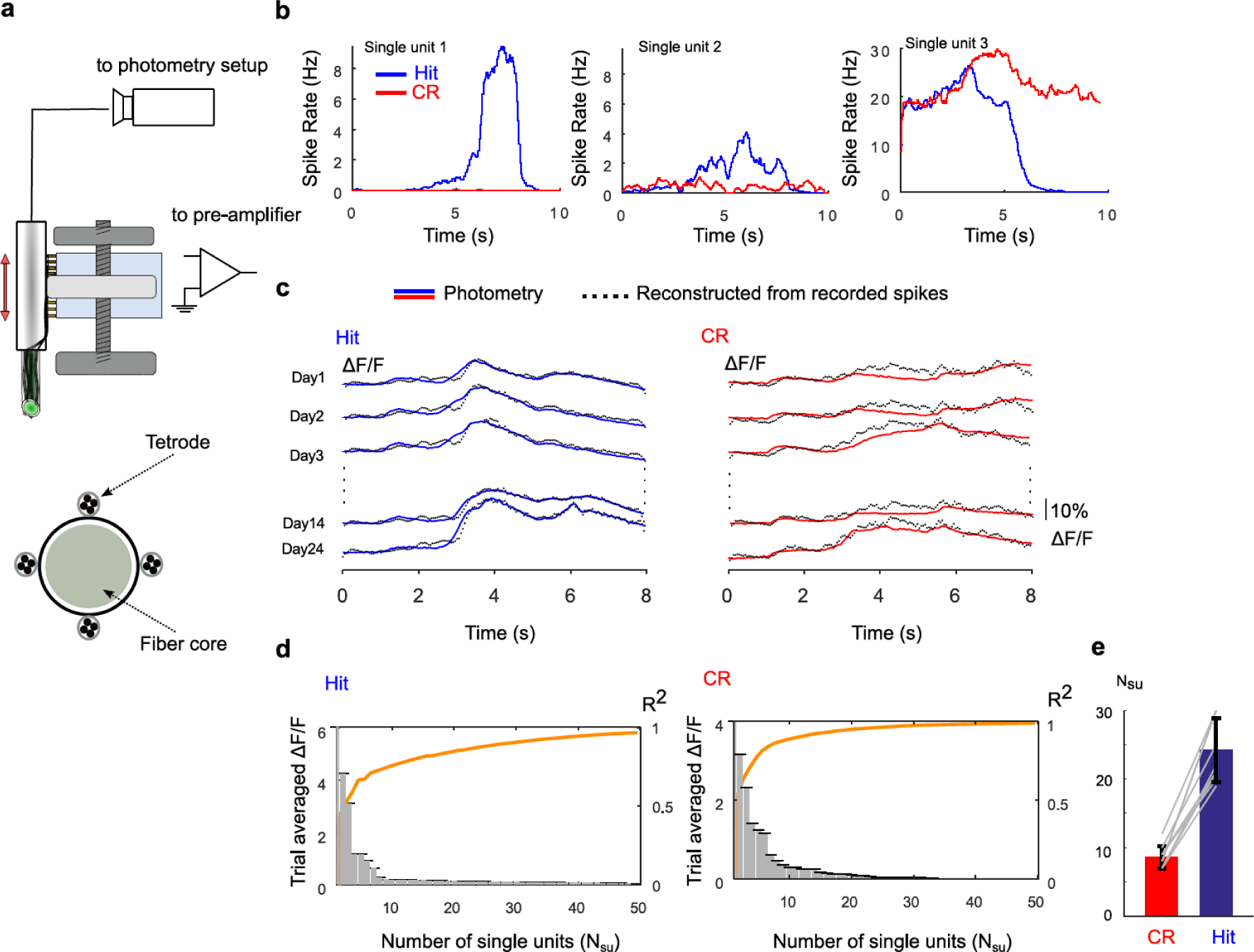
Simultaneous fiber photometry and single-unit recording during texture discrimination. (**a**) Schematic of our custom-designed optotetrode drive for simultaneous fiber photometry and electrophysiological recording. Bottom: Cross section of optotetrode tip comprising the UM22-100 fiber surrounded by four tetrodes. (**b**) Examples of three single units displaying selective spike rate changes in Hit (n = 306) versus CR trials (n = 298). (**c**) Least-squares fit of the fiber-optic bulk calcium signal using the weighted spike rates of 364 single units that were recorded in the vicinity of the fiber tip, convolved with an exponential curve to account for the calcium indicator characteristics. The mean calcium signals for Hit trials (averaged over 187, 257, 251, 306 and 112 trials for day 1 to day 24, respectively) and CR trials (averaged over 181, 227, 237, 298 and 120 CR trials for day 1 to day 24, respectively) are plotted in as blue and red solid traces and the fitted curve is plotted as dashed black trace. (**d**) Bar plots depict the amplitude contribution (in %ΔF/F, left axes) of single units to the fit of the ΔF/F, sorted in descending order. Left: Hit trials; Right: CR trials. The orange solid line shows cumulative R^2^ (explained variance) for the sorted single units. (**e**) Comparison of the number of single units required to explain 90% of the variance in fiber-optic calcium signal for Hit and CR trials (n = 2 mice).

**Supplementary Figure 3.**
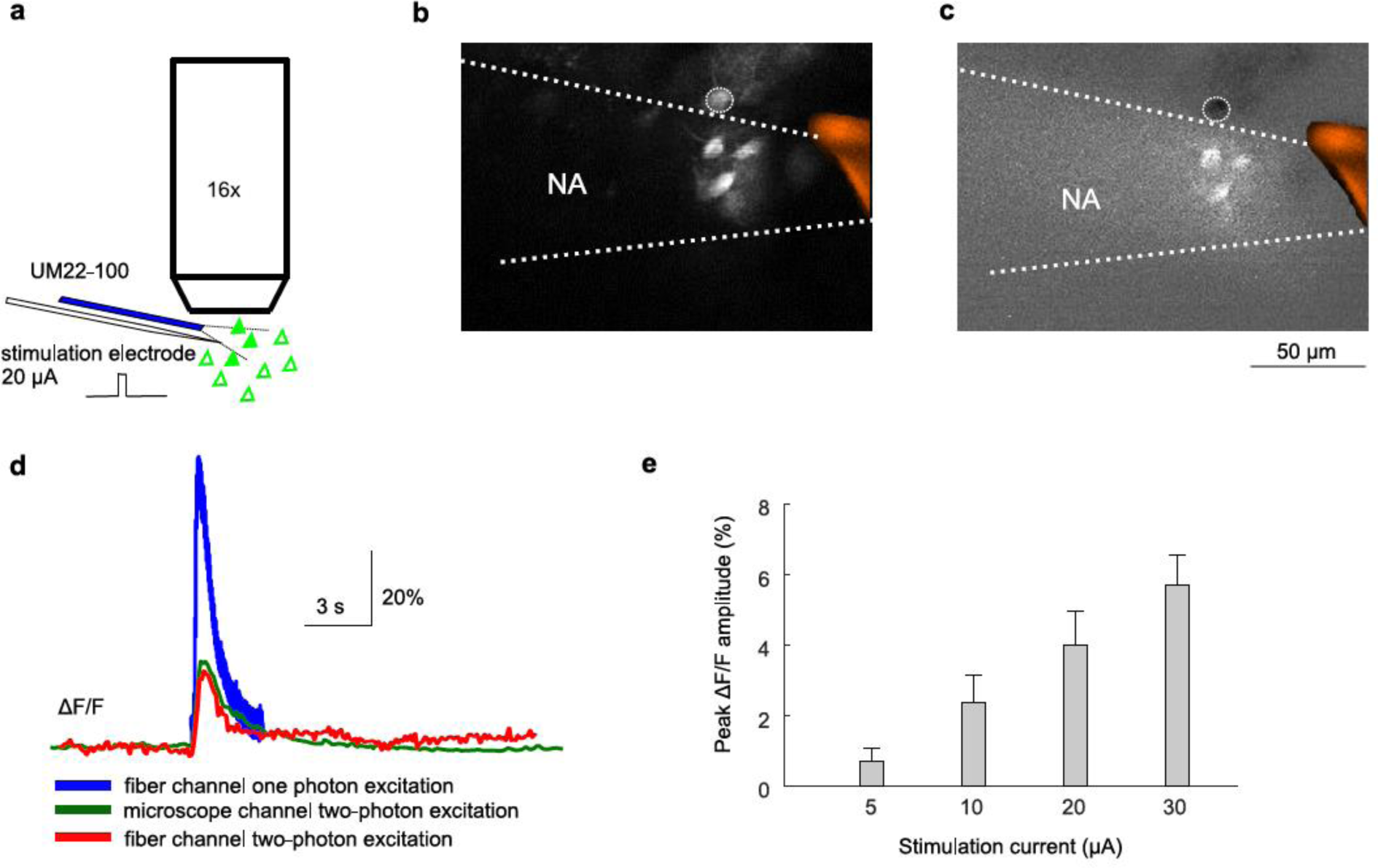
Comparison of fiber photometry and two-photon calcium imaging. (**a**) Schematic of the experiment combining a fiber-optic recording under a 16x microscope objective and two-photon calcium imaging. A piggyback stimulation electrode allowed for local electrical stimulation. (**b**) Two-photon fluorescence image capturing the neuronal ensemble activated by a 50-ms, 20-µA stimulation pulse as detected by the standard photomultiplier in the two-photo microscope detection path The tip of the UM22-100 fiber is shown in red. (**c**) Two-photon fluorescence image upon the same stimulation pulse as collected through the UM22-100 fiber and detected in the photometry setup. Note the neuron that apparently is not ‘seen’ by the fiber tip (marked as circle). (**d**) Comparison of stimulus-evoked ΔF/F calcium transients averaged over the entire field-of-view for the two-photon microscope collected through the objective (green), using scanned two-photon excitation but light collection through the fiber (red), and using single-photon excitation at 473 nm and fluorescence collection by the photometry system (blue). (**e**) Mean ΔF/F amplitudes evoked for stimulation pulses of 5, 10, 20 and 30 µA and measured with fiber photometry.

**Supplementary Figure 4.**
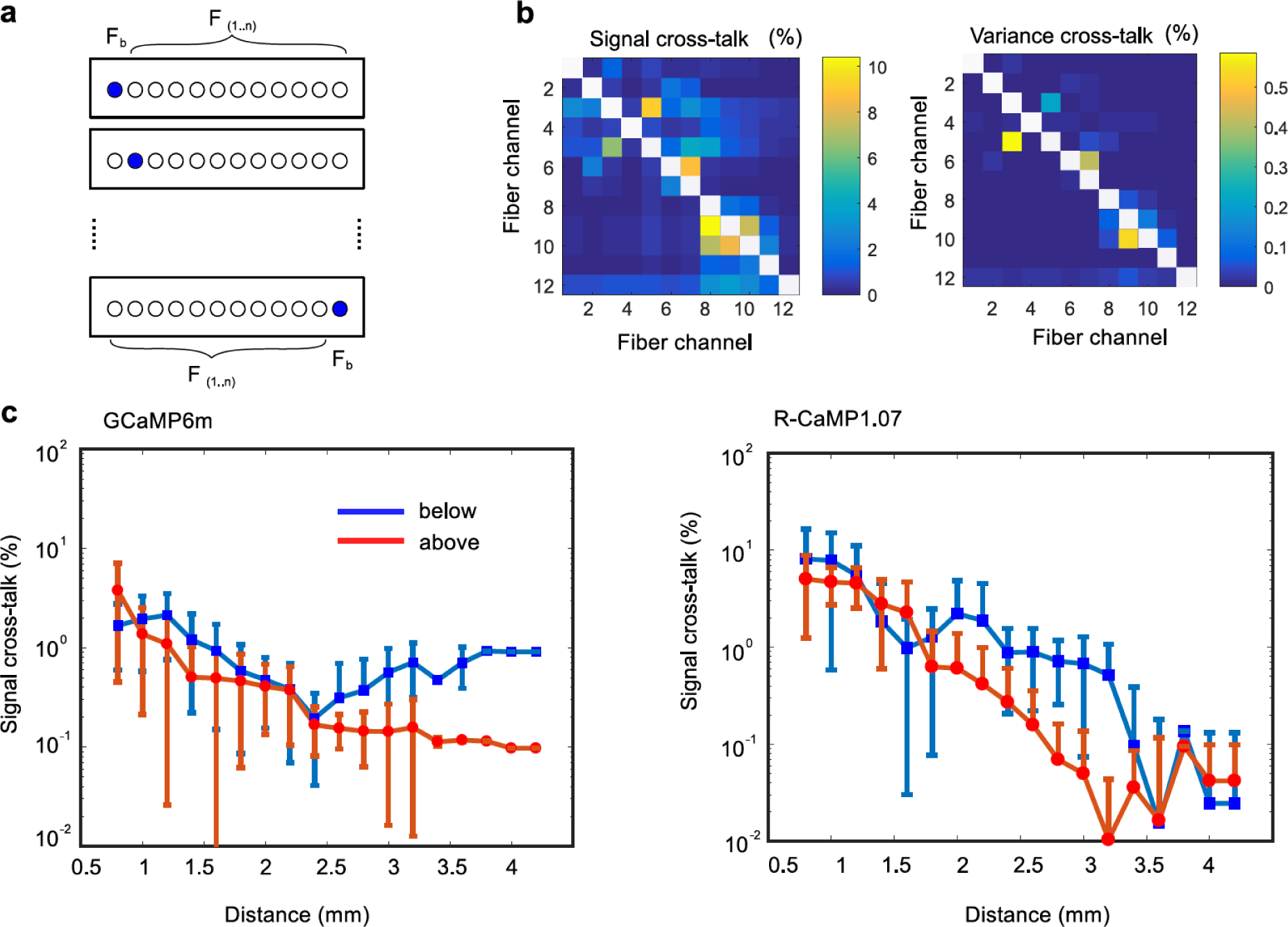
Assessment of potential cross talk between different fiber channels. (**a**) Schematic illustrating the sequential coupling of excitation light into individual fiber channels. To estimate cross talk we coupled excitation light into each fiber separately while measuring fluorescence from all 12 channels. (**b**) Two measures of cross talk. Left: The percentwise contamination was estimated from the median value of the fluorescence recorded in each channel F_1..n_ normalized by F_b_, the median value of fluorescence excited by direct illumination of the respective channel. Right: A second measure is based on evaluating the variance of the fluorescence signal induced by excitation in F_1..n_ normalized by F_b_. (**c**) Median estimation of cross talk for GCaMP6m and R-CaMP1.07 sorted by the distance between the tips of the excited and recorded fiber, shown separately for the cases when the recorded fiber is in front or in the back of the excited fiber.

**Supplementary Figure 5.**
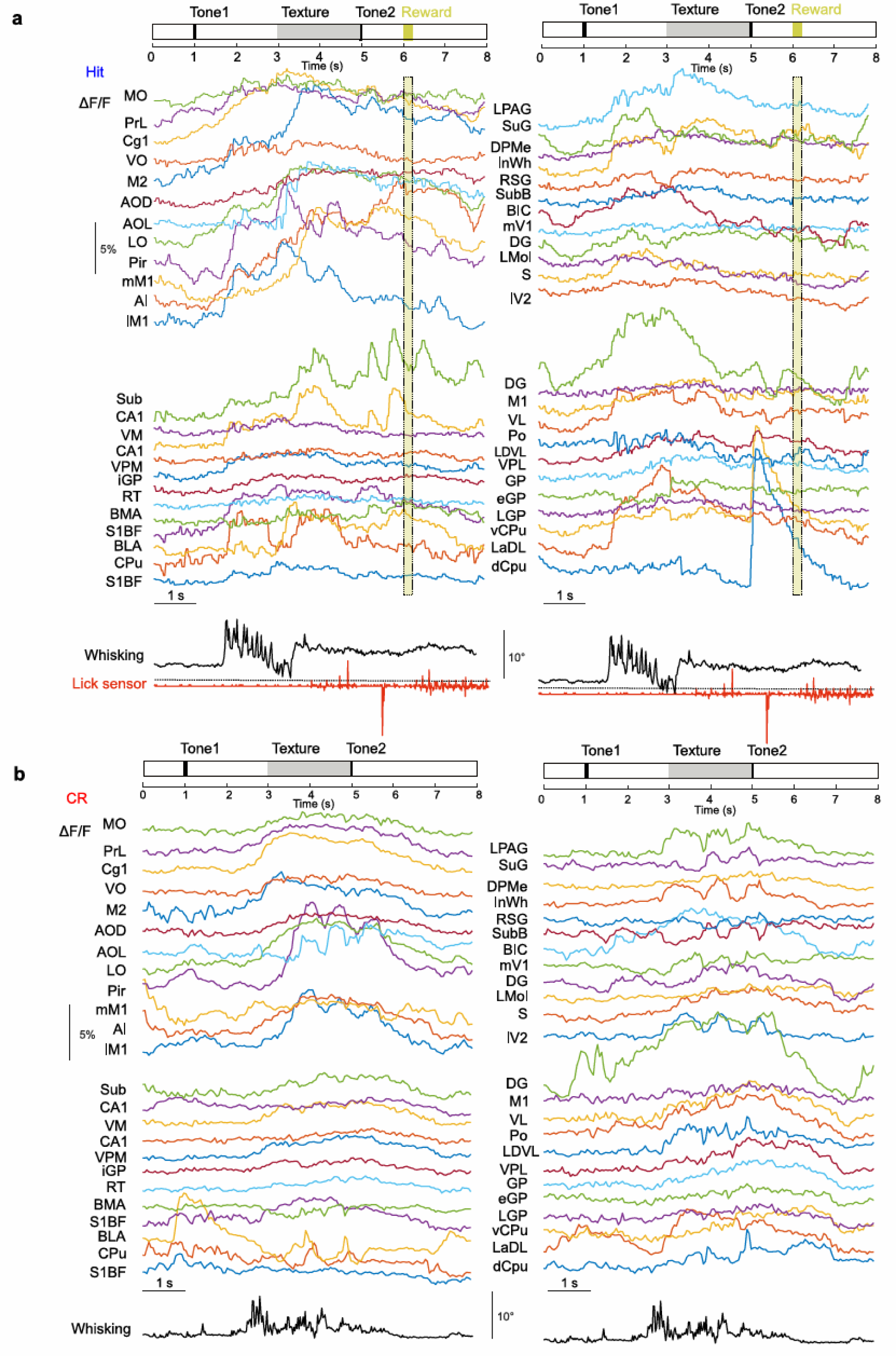
Example of Hit and CR trials for the 48-channel photometry during texture discrimination task. (**a**) Example of a Hit trial. Top: a horizontal bar plot shows a Hit trial structure (black lines indicate time onsets for auditory tones signalling texture approach (starting at 1 seconds), report decision (at 5 seconds); grey bar represents a texture presentation time; yellow bar shows reward period). ΔF/F fluorescence signal is plotted for every channel. Whisking angle and lick events are repeated for each block of 24-channels on the bottom of the inset a and b. (**b**) Example of a CR trial. Top: a horizontal bar plot shows a CR trial (no lick and reward related events).

**Supplementary Figure 6.**
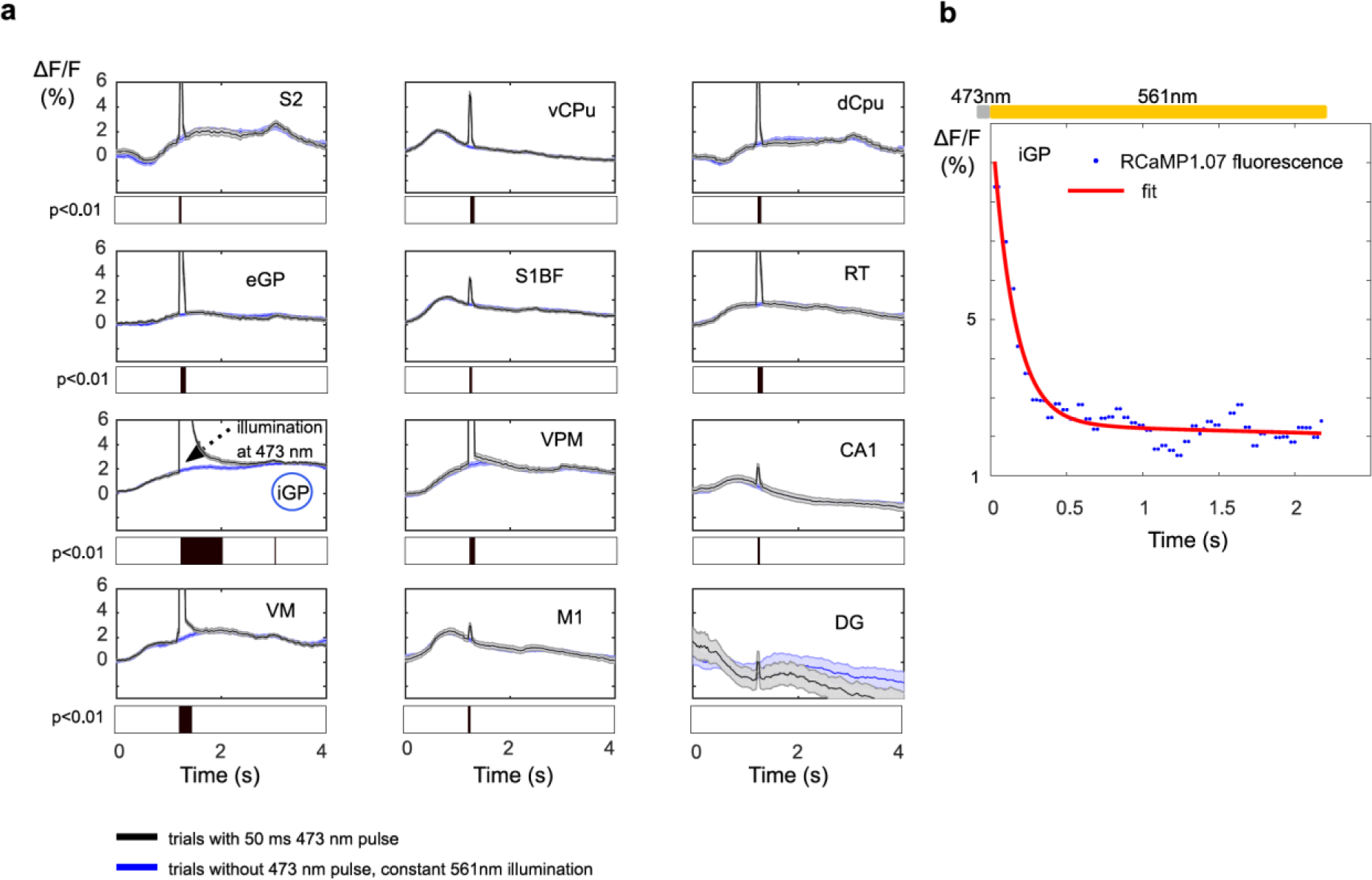
Fluorescence tail after a brief 50ms illumination by 473 nm in C57BL/6 mice expressing only R-CaMP1.07. (**a**) Averaged ΔF/F activity (mean ± SEM) without perturbation laser for Hit trials (n = 134) is plotted with solid blue line. Averaged ΔF/F activity (mean ± SEM) with perturbation laser pulse of 50ms during Hit trials (n = 88) is plotted with dashed black and grey respectively. Both traces were measured at the R-CaMP1.07 emission band. 473 nm laser pulse was coupled to iGP channel only. Bottom (for every channel): black bars indicate significant time samples (p<0.01, one-way ANOVA) for Hit trials with and without perturbation compared. (**b**) Example of a trial with a 473 nm laser pulse and a least squares exponential fit.

**Supplementary Figure 7.**
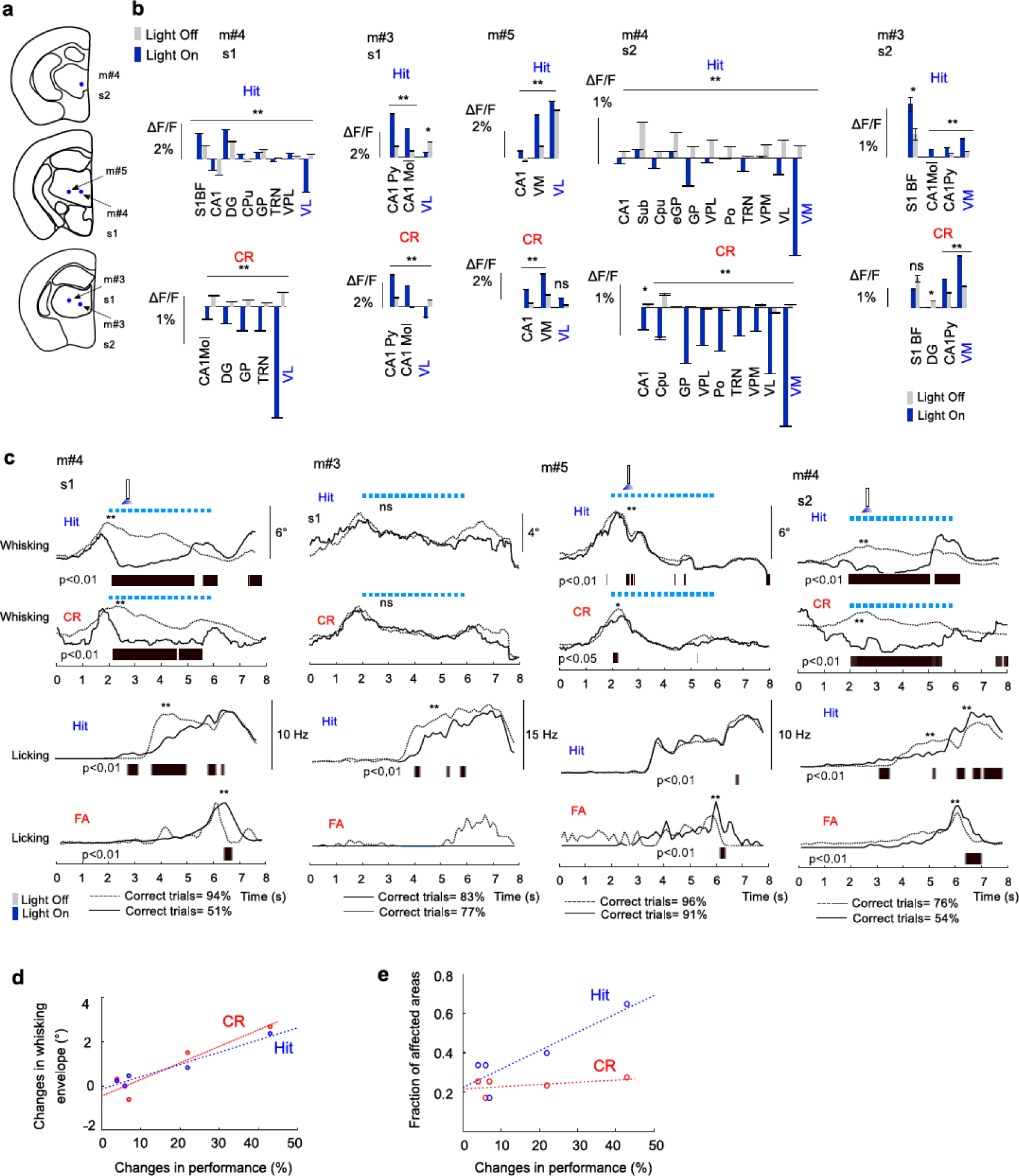
Perturbation of ventral thalamus during a texture discrimination task caused changes in a mesoscale network and behavioral variables. (**a**) Identified fiber tip positions for all mice. Several fibers were implanted into ventral thalamus (m#3 s1 and s2, m#4 s1 and s2 and m#5 s1). Light at 473-nm was coupled into individual fiber. (**b**) Changes in calcium dynamics across recorded channels. Bar plotted in grey for unperturbed trials and blue for perturbed trials. Hit and CR trials are plotted in top and bottom raw respectively. (**c**) Changes in whisking and licking for a full trial duration. Behavioral variables for the perturbed/unperturbed trials are plotted in solid/dashed line respectively. Whisking and licking were reduced during optogenetic perturbation. Mice also developed an impulsive response to a tone signaling for a lick/no-lick i.e. increased lick rate on FA trials (with the exception of m#3). (**d**) Reduction in a whisking envelope during Hit and CR trials correlated to a change in performance. Change in performance is calculated as a difference in a percent of correct trials without/with perturbation. (**e**) Fraction of areas affected by perturbation (number of regions with a significant change in a mean ΔF/F during a post-perturbation period divided by a total number of measured regions) compared to a change in performance.

**Supplementary Table 1.** List of brain regions targeted in the multi-fiber photometry experiments. We mainly adhere to the nomenclature of Paxinos and Franklin (ref. 66). In a few cases we applied further subdivisions, e.g., for medial and lateral M1 and for sublaminae in CA1.

|  |  |  |
| --- | --- | --- |
| 12F anterior | IM1 | Primary motor cortex, lateral part |
|  | AI | Agranular insular cortex |
|  | mM1 | Primary motor cortex, medial part |
|  | Pir | Piriform cortex |
|  | LO | Lateral orbital cortex |
|  | AOL | Anterior olfactory nucleus, lateral part |
|  | AOD | Anterior olfactory nucleus, dorsal part |
|  | M2 | Secondary motor cortex |
|  | VO | Ventral orbital cortex |
|  | Cg1 | Cingulate cortex, area 1 |
|  | PrL | Prelimbic cortex |
|  | MO | Medial orbital cortex |
| 24F medial, or 12F medial (selected subset) | S1BF | Primary somatosensory cortex, barrel field |
|  | S2 | Secondary somatosensory cortex, barrel field |
|  | M1 | Primary motor cortex, posterior part |
|  | CPu | Caudate putamen (striatum), lateral part |
|  | dCpu | Caudate putamen (striatum), dorsal-lateral part |
|  | vCpu | Caudate putamen (striatum), ventral-lateral part |
|  | BLA | Basolateral amygdaloid nucleus, anterior part |
|  | BMA | Basomedial amygdaloid, anterior part |
|  | LaDL | Lateral amygdaloid nucleus, dorsal part |
|  | GP | Medial globus pallidus |
|  | iGP | Globus pallidus, internal part |
|  | eGP | Globus pallidus, external part |
|  | LGP | Lateral globus pallidus |
|  | VPM | Ventral posteromedial thalamic nucleus |
|  | VPL | Ventral posterolateral thalamic nucleus |
|  | VM | Ventromedial thalamic nucleus |
|  | VL | Ventrolateral thalamic nucleus |
|  | LD | Laterodorsal thalamic nucleus |
|  | Po | Posterior thalamic nuclear group |
|  | RT | Reticular thalamic nucleus |
|  | CA1 | CA1 field of the hippocampus, dorsal-lateral lamina |
|  | CA1Py | CA1 field of the hippocampus, pyramidal lamina |
|  | CA1Mol | CA1 field of the hippocampus, stratum lacunosum moleculare |
|  | DG | Dentate gyrus |
| 12F posterior | S | Subiculum |
|  | vDG | Dentate gyrus, ventral part |
|  | vCA1Mol | CA1, stratum lacunosum moleculare ventral part |
|  | mV1 | Primary visual cortex, medial part |
|  | IV2 | Secondary visual cortex, lateral part |
|  | BIC | Nucleus of the brachium inferior colliculus |
|  | SubB | Subbrachial nucleus |
|  | RSG | Retrosplenial granular cortex |
|  | InWh | Intermediate white layer of the superior colliculus |
|  | DpMe | Deep mesencephalic nucleus |
|  | SuG | Superficial gray, superior colliculus |
|  | LPAG | Lateral periaqueductal gray |

